# Individual cell fate and population dynamics revealed by a mathematical model linking telomere length and replicative senescence

**DOI:** 10.1101/2023.11.22.568287

**Authors:** Anaïs Rat, Marie Doumic, Maria Teresa Teixeira, Zhou Xu

**Author notes:** Univ Brest, CNRS UMR 6205, Laboratoire de Mathématiques de Bretagne Atlantique. These authors contributed equally to this work (Marie Doumic, Maria Teresa Teixeira, Zhou Xu).

## Abstract

Progressive shortening of telomeres ultimately causes replicative senescence and is linked with aging and tumor suppression. Studying the intricate link between telomere shortening and senescence at the molecular level and its population-scale effects over time is challenging with current approaches but crucial for understanding behavior at the organ or tissue level. In this study, we developed a mathematical model for telomere shortening and the onset of replicative senescence using data from *Saccharomyces cerevisiae* without telomerase. Our model tracks individual cell states, their telomere length dynamics, and lifespan over time, revealing selection forces within a population. We discovered that both cell genealogy and global telomere length distribution are key to determine the population proliferation capacity. We also discovered that cell growth defects unrelated to telomeres also affect subsequent proliferation and may act as confounding variables in replicative senescence assays. Overall, while there is a deterministic limit for the shortest telomere length, the stochastic occurrence of non-terminal arrests drive cells into a totally different regime, which may promote genome instability and senescence escape. Our results offer a comprehensive framework for investigating the implications of telomere length on human diseases.

**One-sentence Summary:** Key determinants of population proliferation capacity in the context of telomere shortening.

## Introduction

Mathematical modeling plays a pivotal role in elucidating the underlying principles and mechanisms governing biological processes. However, developing models that capture the heterogeneity of biological systems is challenging, particularly when heterogeneity evolves over time during population growth. This occurs during tumorigenesis, for example, where increasing genomic instability can affect cell morphology, proliferation rates, invasion potential and resistance mechanisms (*1*). Another example is the phenotypic heterogeneity that accumulates over time in the course of replicative senescence caused by telomere shortening (*2*).

In human somatic cells, telomere shortening eventually activates the DNA damage checkpoint, arrests the cell cycle and triggers replicative senescence, which is thought to contribute to human aging (*3*), (*4*). The loss of telomere length homeostasis is also a characteristic of cancer cells (*5*). Likewise, abnormally short telomeres are responsible for a spectrum of deadly telomeropathies, which currently remain incurable (*6*). However, the mechanisms linking telomere shortening with replicative senescence at the population level remain poorly understood (*2, 7-11*).

Phenotypic heterogeneity in replicative senescence cannot be deciphered in population-based experiments, in which senescence appears as homogeneously progressive. In contrast, single-cell analyses, mainly obtained in telomerase-negative *Saccharomyces cerevisiae*, have revealed that senescence of individual telomerase-negative cells is highly variable (*12-14*). In addition, senescence onset is often abrupt: cells switch in a single division from fast proliferation to a prolonged cell cycle, which can be followed by a few more long cell cycles before cell death. Below we use the term replicative senescence or senescence to refer to this ultimate sequence of long cell cycles, qualified as terminal since they lead to cell death in the budding yeast model.

Because the onset of senescence may happen at different times, even for descendants of the same cell, it is an intrinsically asynchronous process. Numerical and experimental evidence in *S. cerevisiae* support a model in which senescence onset is triggered by the shortest telomere reaching a threshold length, from which the DNA damage checkpoint is first activated (*11, 15-18*). Alternative hypotheses proposing that replicative senescence could be triggered by multiple telomeres or simply by the passage of time fail to explain the substantial variability observed in the onset of senescence. Therefore, a significant contributor to asynchrony arises from the shortening trajectory of the shortest telomeres within a cell. This trajectory is driven by the process of telomere replication, which inherently generates asymmetry and variability, as observed at the molecular level in *S. cerevisiae* and other organisms (*19-22*).

In addition to the variability in senescence onset, another source of heterogeneity during the onset of senescence was observed in single telomerase-negative budding yeast cells (*13, 14*). Specifically, a subset of individual cells, so-called *type B* cells, undergo several switches between non-terminal arrests and normal proliferation prior to terminal senescence, in contrast to *type A* cells, which undergo the switch in a single step. Even though both terminal and non-terminal arrests are abnormally long cycles associated with the activation of the DNA damage checkpoint, their manifestation corresponds to distinct probability laws, thus possibly arising from distinct molecular origins (*23, 24*). In non-terminal arrests of *type B* cells, the cells either recover upon repair of a damage, presumably stemming from a yet unidentified telomere defect, or adapt to the damage, forcing mitosis until successful repair. These cycles of mitosis in the presence of unrepaired DNA damage are a source of genomic instability, which likely fuels additional heterogeneity.

While these single-cell level observations and population experiments that determine the average or use only single time-points have provided insights up to a point, these two types of information would need to be combined to decipher the key determinants of onset of replicative senescence *in vivo*. For instance, the generational age or length of the shortest telomere triggering replicative senescence cannot currently be accessed experimentally in populations or individual cells. Thus, to study specific *in vivo* dynamics requires modeling senescence at the population scale. This provides a biologically-relevant scale for understanding tissue development and renewal, aging, cancer emergence and growth. However, this is challenging for several reasons, including the complex effects of competition and selection, and the poorly understood effects of telomere shortening, cell death, terminal or non-terminal arrests on population dynamics.

To overcome these experimental limitations, we built a mathematical model of telomerase-negative cell population growth, and used single-cell data to calibrate it. We used *S. cerevisiae*, in which telomerase can be inactivated, as a well-documented experimental model of replicative senescence (*25-27*). We thus exploited the wealth of information and molecular models derived from *S. cerevisiae* regarding telomere shortening pathways, the molecular mechanism underlying replicative senescence, and the data from single-cell lineages in microfluidics experiments (*11, 14, 16, 18, 23, 24*), to track in depth the state of each individual cell of the population, including the length of each telomere over time. We managed to delineate hidden components of replicative senescence mechanism beyond the scope of experimental observations. Specifically, we found an almost deterministic limit of telomere length triggering senescence, existing alongside a different regime with a fully stochastic route to senescence, driven by the random occurrence of non-terminal arrests. This route, together with the strong selection of cells carrying the longer shortest telomeres in the initial population, offers potential senescence escape. Our results also enable an assessment of the consequences of other independent comorbidities on lifespan with consequences for our understanding of age-related human diseases.

## Results

### Rationale of the mathematical model

We built a population model and used *in silico* experiments to investigate the structure and evolution in time of a population of yeast cells in which telomerase is inactivated. In order to validate the model with experimental data, our strategy was to mimic actual senescence experiments in which population doublings are monitored each day from telomerase inactivation to senescence crisis, i.e. when the culture reaches the lowest growth capacity, typically a few days after telomerase inactivation. In order to focus exclusively on the first driving forces of senescence mechanisms, and because of the lack of data at the single-cell level, our model does not take into account the post-senescence survivors, whose frequency was globally estimated at ∼2.10^−5^ (*28*). Because cells grow very robustly and exhaust nutrients during the first days of the experiment, cultures are diluted each day in fresh media, so only a fraction of the population is sampled for the next day. In this assay, the cell concentration and the statistical mode of telomere lengths, i.e. the peak of the telomere length distribution, can be measured each day (Fig. 1A).

**Figure 1.**
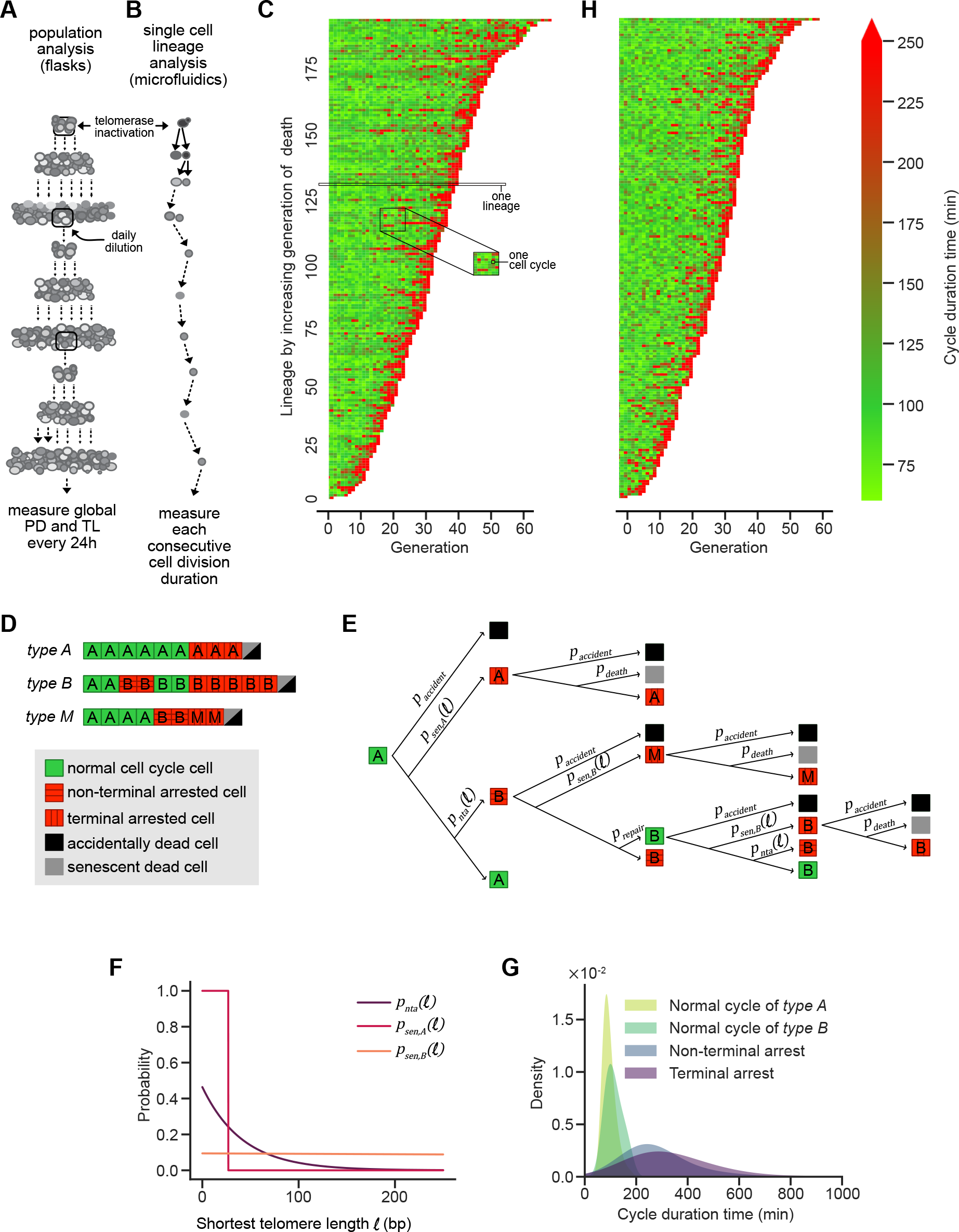
Principle and description of the mathematical model of replicative senescence. (A-B) Schematic representation of population (A) and single-cell lineage (B) experiments. (C) Experimental data of microfluidics experiments from (*23*). Display of consecutive cell cycle durations of TetO2-TLC1 lineages. Generation 0 corresponds to the moment from which telomerase is inactivated. Each horizontal line is an individual cell lineage, and each segment is a cell cycle. Cell cycle duration (in minutes) is indicated by the color bar on the right. (D) Three examples of lineages containing indicated *type* cells (*A, B* or *M*). The colors of the squares indicate whether the cell cycle is normal (green) or abnormally long (red). Vertical or horizontal strips specify whether the cell cycles are terminal (never followed by a normal cell cycle) or not, respectively. Black or gray cells indicate the type of death. (E) Tree diagram of the mathematical model. It indicates the several fates of each cell *type* and respective probability rates, some being functions of the length of the shortest telomere in the cell *ℓ*. Legends as in (D). (F) Best fits of indicated probability rates as functions of the length of the shortest telomere *ℓ*. (G) Probability densities of cell cycle durations for indicated cell types. (H) Simulated data of one microfluidics experiment with the best-fit mathematical model (parameters in Table 1). Other examples are shown in Fig. S1F-G.

This typical population experimental design introduces several biases. Firstly, cell-to-cell variability in cell cycle duration times introduces competition (fast-dividing cells rapidly take over the population). Secondly, senescence-induced death also introduces selection effects: fast-dividing cells, although over-represented initially, are likely to die first because they reach their proliferative limit faster in time. In contrast, slow-dividing cells can take over the population in the long term. Therefore, in a replicative senescence culture, the number of population doublings is not linearly related to the number of generations undergone by cells, and consequently, daily telomere length measurements cannot be directly used to estimate a shortening rate per generation. Besides, daily dilution constitutes another sampling bias.

**Table 1:** Parameters fitted on the microfluidic data and used for the population simulation experiments

| parameter | value |
| --- | --- |
| $a_{nta}$ | 0.02 |
| $b_{nta}$ | 0.44 |
| $a_{sen,A}$ | 0.19 |
| $b_{sen,A}$ | 0.73 |
| $a_{sen,B}$ | 0 |
| $b_{sen,B}$ | 0.12 |
| $\ell_{min,A}$ | 27 |
| $\ell_{min,B}$ | 0 |
| $\ell_{trans}$ | 0 |
| $\ell_0$ | 40 |
| $\ell_1$ | 58 |

The mathematical model is largely based on our data from single cell lineages observed in microfluidic experiments and accounts for a number of already well-characterized types of cell-to-cell variability, as described in (*16, 18, 24*). In the microfluidic experiments, the consecutive cell cycle durations are measured from telomerase inactivation to cell death in individual cell lineages (Fig.1B-C). These quantitative data enabled us to define senescence pathways and their estimated frequency, i.e. the presence or absence of non-terminal arrests, and the switch from *type A* to *type B* cells. *Type B* cells may senesce during the course of their first consecutive non-terminal arrests. These arrests cannot be experimentally distinguished from *type A* senescence arrests. We thus introduced in our model a “misclassified” *type M*, corresponding to *type B* lineages which would have been experimentally labeled as *type A* (Fig. 1D-E). Hereafter we will often refer to any cell that falls under the category of *type B* or *M*, i.e., cells whose ancestors or the cells themselves have experienced a non-terminal arrest, as *type B* cells. Though important for calibrating parameters, this misclassification appears not to change the results qualitatively.

The model requires a detailed description of the population at the single cell level: each cell is tracked in time through its state *S* = (*L, T, C, τ, X*), namely its:

- telomere lengths *L* ∈ *Z*^2*x*16^ (there are 16x2=32 in a haploid *S. cerevisiae* cell which contains 16 chromosomes)
- cell type *T* ∈ {*A, B, M*}
- cell cycle type *C* ∈ {*nor, nta, sen*}, for normal duration cell cycle (*nor*), or prolonged cell cycle, which includes non-terminal arrest (*nta*) or terminal (senescence) arrest (*sen*), respectively,
- cell cycle duration time *τ* ∈ *R*_+_
- and a set *X*of individual variables of interest (generation, birth time, ancestor index in the initial population, etc.)

This description of the population thus goes beyond the accessible population data.

### Initialization and progression of the model

We started with *N*_*init*_ non-arrested cells of *type A* and generation 0. The initial 32 telomere lengths of each cell were drawn randomly and independently, from the distribution described in (*11, 16*); minor modifications on this distribution are described in the Methods section and (Fig. S1A).

We let cells evolve by consecutive cell divisions. At each cell division, transition probabilities from a given mother cell’s state to its daughters rely on the three following submodels: (i) The telomere shortening model (*18*); (ii) Laws of arrests / cell cycle types transition (Fig. 1D-F); and (iii) Laws of cell cycle duration times (Fig. 1G). The length of the shortest telomere in cells was computed after running the telomere shortening model (i) described in (*18*). The transition laws (ii) are driven by the cell and cell cycle types together with the length of the shortest telomere in the cell and account for the two regimes described in (*14, 24*) (Fig. 1D-F). We built upon previous work (*18, 24*) to estimate the stochastic laws describing the entry in a sequence of arrests and exit by either repair or death for non-terminal or terminal arrests, respectively. To these laws, we added a probability of spontaneous death, assumed to occur at a constant rate. These events were assumed to happen at birth (independently in each daughter cell) as summarized in Fig. 1E. The cell cycle duration of each cell (iii) was then drawn from the experimental distribution corresponding to its cell type (Fig. 1C, G).

Every 24 hours, the whole population was diluted: we sampled, randomly with equiprobability, a number *N*_*dil*_ of cells (*N*_*dil*_ = *N*_*init*_) to initiate the next daily culture. As soon as the population reached a saturation number *N*_*sat*_, we stopped making cells divide until the next dilution.

### Calibration of the model

The laws of arrests 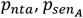 and 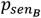 (Fig. 1F) were fitted altogether on the basis of the generations of arrest from the microfluidics data, using the CMA-ES optimization algorithm (Fig. S1B-D). The probabilities to exit a sequence of arrests (*p*_*repair*_ and *p*_*death*_) were assumed to be constant, as demonstrated (*24*). For senescence onset, 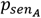 was found to be nearly deterministic and 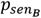 independent of telomere length (Fig 1F). This corresponds to two clearly distinct pathways to senescence. A cell can thus follow two scenarios:

1. Divide for a number of generations of normal cycles until one telomere reaches the almost deterministic threshold 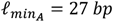. This is the canonical pathway of *type A* lineages, which confirms the deterministic model of (*16*), which focused exclusively on senescence onset of *type A* lineages;
2. Or experience at least one non-terminal arrest to become *type B* with probability *p*_*nta*_. Then, the cell has a constant probability of entering senescence at each cell division, which results in a great variability in the length of the telomere triggering senescence of *type B* (possibly longer or much shorter than 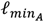).

With these laws calibrated (Table 1), we next simulated the evolution of cell lineages as grown and analyzed in the microfluidics experimental setting (Fig. 1H, S1E-G). The obtained profile was remarkably similar to the experimental one (Fig. 1C). By considering the cumulative simulated data for 1000 independent virtual experiments (Fig. S1D-E), we were able to quantitatively compare the experimental data with the simulated one. The median and variance of experimental lifespan of lineages fit very well within the confidence interval values obtained in the simulated data (Fig. S1D). Likewise, the simulated percentage of *type B* cells lies between 61% and 89%, consistent with the range of experimental *type B* proportion of 61-67% (see Supplementary Material). Altogether, our mathematical model correctly recapitulates the experimental microfluidics data.

### Mathematical model of a senescent population

We next wished to simulate a senescence experiment performed in a population (Fig. 1A, S2A-B). Different laboratories use variations of the same protocol. In the experimental example we used, cultures starting at an optical density measured at 600 nm (OD_600 nm_ or OD) of 0.0125 are grown exponentially in rich media. Wild-type strains are expected to reach 9-9.5 OD, which corresponds to ∼9.5 cell divisions (Fig. S2A) (*20, 29*). Since wild-type budding yeast divides in 90 min on average in rich media (Fig. 1G), this means that saturation is reached after ∼15 h. The culture is then diluted every 24 h in fresh media back to OD = 0.0125.

Simulating the exponential growth of such a high number of cells (saturation corresponds to up to 3.10^8^ cells/ml) is too expensive in terms of computation and allocated memory to keep track of all the parameters of the cells (Fig. S2B). We thus tested whether simulating smaller populations generated a bias. To this aim, we identified a minimal size *N*_*init*_ = 300 at which populations were big enough to account for the experimental data in terms of cell growth (Methods and Extended Materials & Methods). We then defined a saturation ratio *r*_*sat*_ in order to stop the simulation each day when the number of cells reaches *N*_*sat*_ = *r*_*sat*_ *N*_*init*_.

### Validation of the model with data extracted from population senescence assays

Based on the above, we simulated cultures of telomerase-negative cells (Fig. 2A). To test validity, simulated curves were compared with experimental population data, specifically the daily monitoring of the cell concentration and the mode of telomere length distribution extracted from (*20*) (Fig. 2B-C). In this setting, since saturation is reached after ∼9.5 divisions, we set r_sat_ = 720 ≈ 2^9.5^. We observed in Fig. 2B that simulations of the number of cells and experimental measurement of OD were in good agreement for the first 4 days, before there was a drop in proliferation capacity starting at day 5 in both experimental and simulation curves. The same applied for telomere length, in which experimental and simulated telomere shortening fit well, though discrepancy started at day 3. The curves of the experimental growth and telomere length profiles were maintained above the simulated profiles from days 4-5 and after. This suggests that our model recapitulates well the events at work in the majority of cells present in the population during the first days spent in the absence of telomerase. A reason explaining the difference between the experiment and the simulation is that we started the simulation from 300 cells instead of∼3.10^5^, so that extreme cases have less chance to be part of the initial set. This has two consequences. The first is that we omitted post-senescent cells and their descendants, which appear at a low frequency of ∼2.10^−5^ (*28*). Post-senescent survivors are expected to contribute to the increase of cell growth and telomere length, but when they initially appear is currently unknown. Based on the comparison between our simulations and experimental data, they may be already present at day 3 or 4 in a non-negligible proportion. The second consequence is that some extreme cases, like cells having very long initial telomeres, would survive much longer than others and would then greatly contribute to the population.

**Figure 2.**
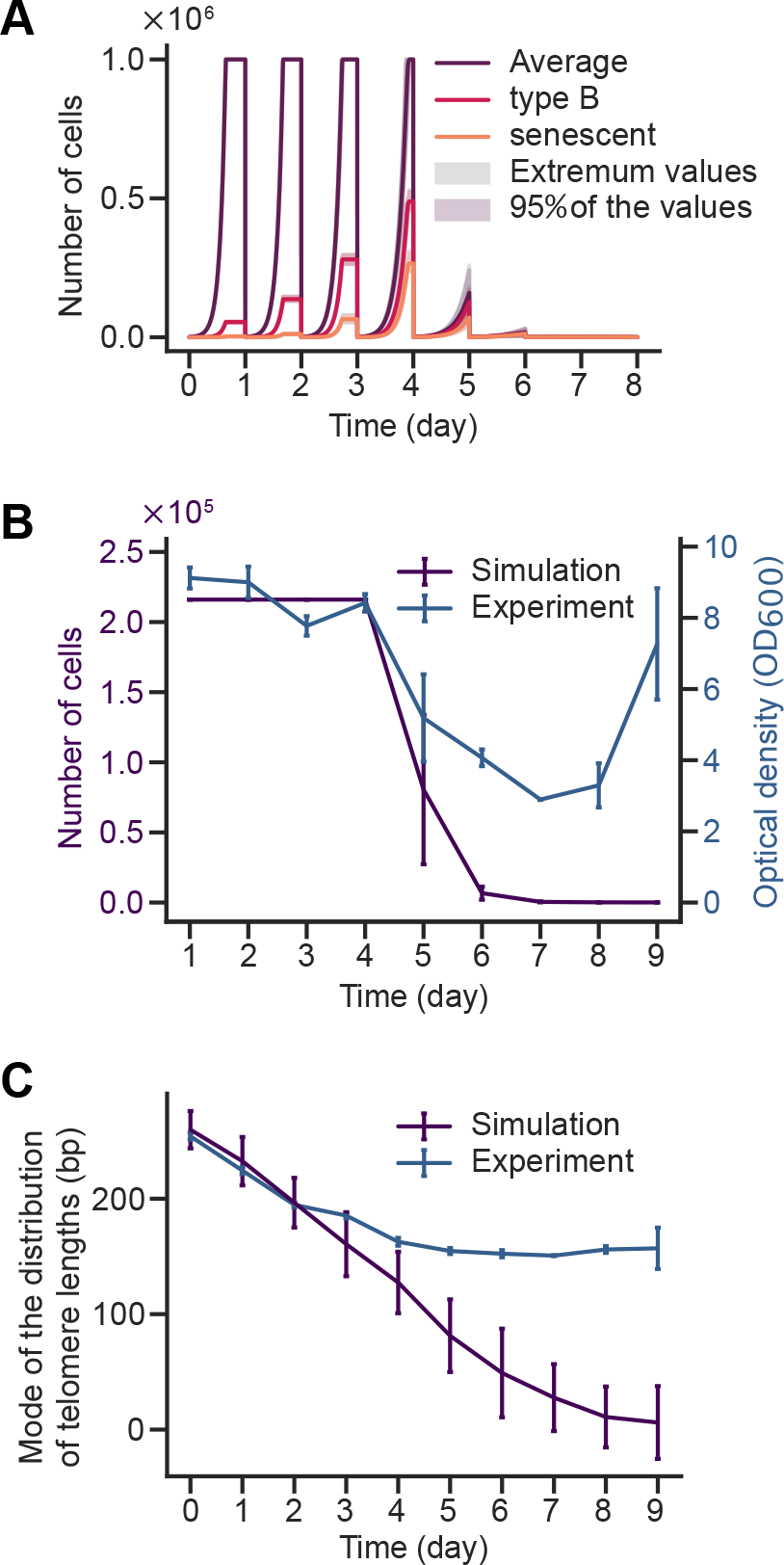
Mathematical model applied to population experiments. (A) Simulation of population growth as a function of time, as schematized in Figure 1A, with best-fit parameters of Table 1 and each day an initial number of cells *N*_*init*_ = 1000, reaching a saturation number *N*_*sat*_ = *r*_*sat*_ *N*_*init*_, with *r*_*sat*_ = 1000. (B) Comparison between experimental data from (*20*) of telomerase-negative cells, and simulations using the best-fit model with *N*_*init*_ = 300 and *r*_*sat*_ = 720, corresponding to the experimental conditions (dilution starting at *OD*^600^ = 0.0125 and reaching saturation at *OD*_600_≈ 9 after 24h of growth in telomerase-positive conditions (see Figure S2A). Only simulated values at the experimental times of observation are displayed, once per day. Error bars of experimental values correspond to SD of 3 independent experiments. Error bars of simulated data correspond to SD of 30 independent simulations. (C) Comparison of the telomere length mode between the same experiments and simulations as in (B). Error bars of experimental values correspond to SD of 3 independent experiments. Error bars of simulated data correspond to SD of 30 independent simulations.

Another reason that might explain the difference lies in the method used to experimentally inactivate telomerase (the TetO2 repressible promoter): it might display some leakage, so that a few critically short telomeres could be specifically elongated by very few active telomerase molecules in cells (*30*). Similarly, although this may apply to only very few cells, they may become dominant over time when all other cell descendants have entered senescence.

Having these two bias sources in mind, we concluded that our simulations best recapitulate the dynamics of a population in the absence of telomerase within the first 3-5 days after loss of telomerase activity. Because it omits the post-senescence survival pathway, our model should help dissect the causes and consequences of the mechanisms contributing to the decrease of cell proliferation (i.e., replicative senescence) in the absence of telomerase.

While varying *r*_*sat*_ had some effect on senescence kinetics (Fig. S2C-D), for the rest of the study, we rounded the number of population doublings to reach saturation to 10 (*r*_*sat*_ = 1000).

### Assessing quantitative data on population structure and evolution

Our simulations also enabled quantification of the heterogeneity of replicative senescence cultures in terms of composition of cell “age”, expressed in generations. As time progressed, the number of generations undergone by cells increased rapidly in early cultures and more slowly in late cultures (Fig. 3A). This reflects the fact that cells that have undergone prolonged cell cycles remain longer in the cultures, so that there is a progressive replacement of *type A* cells by *type B* cells (Fig. 3B). The variance of these generations also increased substantially with time, reflecting the increasing heterogeneity of cultures. Notably, the proportion of senescent cells became substantial prior to the experimentally measurable decline of cell proliferation in cultures (Fig. 3C). At days 3-4, while one cannot detect a decline in the population proliferation potential (Fig. 2B), 10-25 % of the cells in the cultures are actually senescent. When we detailed the composition of senescent cells according to cell *types* (*A* or *B*) (Fig 3C), we observed that on the first day, the vast majority of the cells entering senescence were *type B*, partially misclassified as *type A*, since telomeres of *type A* cells have not yet approached the threshold 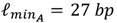. On day 2, *type A* cells started to enter senescence and from day 3 to day 5, there were significantly more *type A* cells entering senescence than *type B*. The last cells entering senescence from day 5 onward were mostly *type B*. We concluded that tracking the global biochemical properties of aging populations over time inadequately reflects the individual cell-level changes. Instead, it indicates a turnover of senescent cells following various trajectories from the earliest time points.

**Figure 3.**
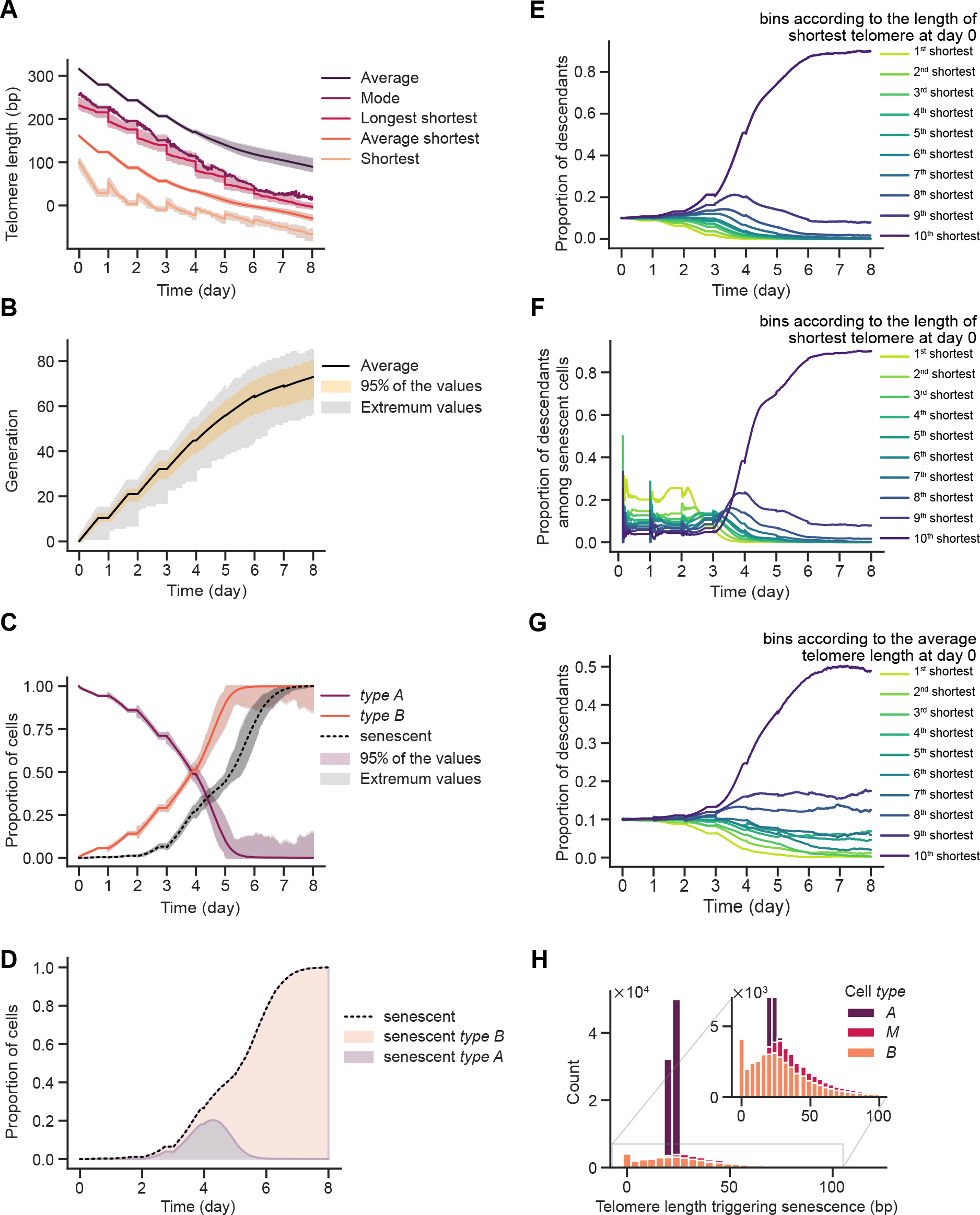
Hidden parameters, experimentally inaccessible, extracted from simulations of telomerase-negative cell cultures. (A) Telomere length of indicated telomere features. (B) Generational age distribution evolution with time. (C) Evolution over time of the proportions of indicated cell categories. (D) Evolution over time of the proportion of senescent cells in the total population (black dotted curve) and the proportion of type A or type B cells among the senescent population. (E) Proportion of descendants according to the initial shortest telomere length of their ancestors. (F) Same as (E) for senescent cells only. (G) Proportion of descendants according to the average telomere length of their ancestors. (H) Distribution of the length of the shortest telomere in senescent cells for indicated cell *types*.

Given the ongoing debate regarding whether average telomere length or average shortest telomere length serves as the most accurate proxy for biological age (*31*), we next wished to use our simulations to estimate values that cannot be measured experimentally and evaluate this issue. We plotted the average and the mode of telomere length distribution in the population over time, and the evolution of the shortest telomere (represented by the average shortest, the shortest shortest, or the longest shortest) in the cell in the absence of telomerase (Fig. 3D). We found that all telomere length values shorten in a nonlinear fashion, faster in the first days upon telomerase inactivation, compared to late cultures. This is due to the continuous selection of fast-growing cells possessing longer telomeres. This was more prominent for the shortest telomere in cells, since it is the telomere to which the stronger selection applies. We concluded that while the length of the shortest telomere is the major determinant of the onset of senescence, the telomere length mode, as well as the average of the length of the shortest telomeres, can effectively act as a nonlinear but reliable proxy for the age of a population. Notably, this somewhat counterintuitive effect is anticipated to be amplified as the initial population exhibits a greater degree of variation in telomere lengths among its shortest telomeres, a characteristic linked to the genealogy of the initial cells and the mechanism of telomere length homeostasis prior to telomerase inactivation.

To illustrate further this effect on the shortest telomeres, we also studied how the population is structured over time as a function of the length of the shortest telomere in the initial ancestral cells. To this end, we clustered the initial cells into 10 equally sized bins according to the length of their shortest telomere, and recorded the number of cells derived from each of these initial cells over time (Fig. 3E-G). We observed that starting at day 3-4, an increasing proportion of the population stemmed from a single population of cells with the longest shortest telomeres, reflecting the selection based on the length of the initial shortest telomere in the cell. Conversely, the progeny of the cells with shorter shortest telomeres constituted the first senescent cells found in cultures, while the cells having longer shortest telomeres entered senescence later (Fig. 3F). In contrast, as expected, ordering the initial cells by increasing average telomere length revealed a lesser selection effect (Fig. 3G). Among *type B* cells, we found an early selection effect for cells displaying shorter shortest telomere, meaning that although they undergo a non-terminal arrest early in culture, cells may preserve a significant proliferation potential, a tendency that disappears rapidly (Fig. S3A).

Lastly, we estimated the length of the shortest telomere in cells when entering senescence. We found as expected that a large fraction of cells (*type A*) entered senescence when the shortest telomere was around 27 bp, which is the 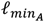 value (Fig. 3H and S3B). Unexpectedly, for *type B* cells (and *type M*), the length of the shortest telomere was widely distributed, ranging from 0 to 70 bp (comprising ∼95% of the values). This reveals variability introduced by a first non-terminal arrest: it drives cells into a totally different regime and a much more variable route to senescence.

### Influence of initial telomere length distribution and mortality on population dynamics

Many mutants and growth conditions are known to affect telomere length homeostasis, and are expected to alter senescence rates (*32-36*). To quantify the consequences of altered initial telomere length distribution on the telomerase-negative cell population, we ran simulations of populations in which the initial telomere length distribution was modified by a translation of the whole telomere length distribution (see Methods). To estimate the effect on senescence rate in a cell population, we defined the time when only half of the saturation limit is reached (HSL). We found that HSL varied with the global translation of telomere length distribution in a non-linear way. A translation of -20 bp or +20 bp in telomere length homeostasis led to a change of -14 h or +17 h in HSL, respectively (Fig. 4A and S4A). As expected, average telomere length was shorter when initial telomere length was shorter and the shortening rate was maintained for the first 4 days (Fig. 4B and S4B). We then altered telomere length distribution by dilatation of the left side of the distribution by *ℓ*_0_ and, unexpectedly, observed only minor effects on the HSL (Fig. 4C and S4C). This counterintuitive result can be explained, again, by the pronounced selection pressure acting on cells with longer shortest telomeres, under the condition that the initial population size and telomere length heterogeneity is sufficient. Because we altered only the distribution of the shortest telomeres, the average telomere shortening remained mostly unchanged (Fig. 4D and S4D).

**Figure 4.**
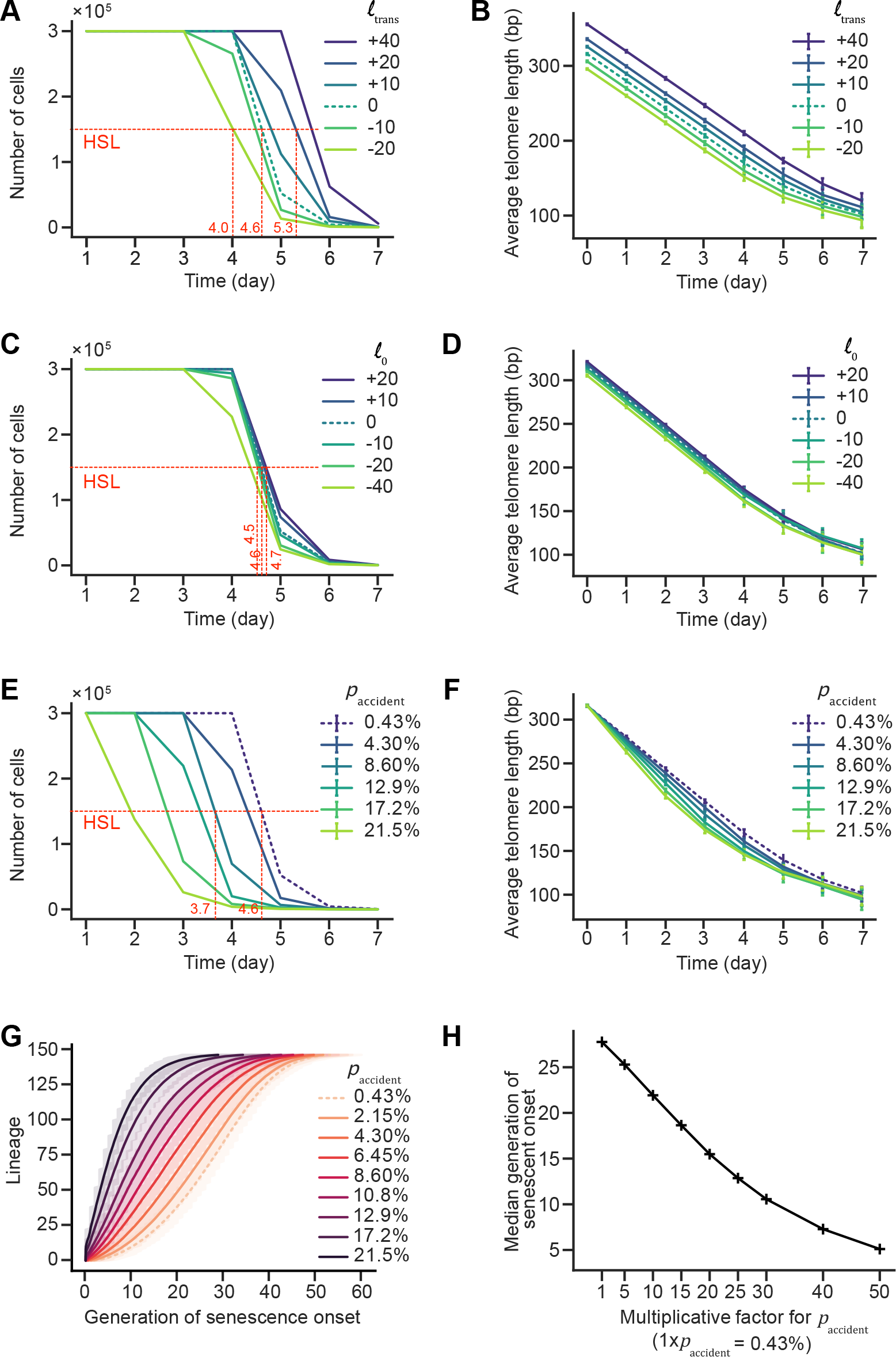
Effects of altering telomere length prior to telomerase inactivation or the constant mortality rate on replicative senescence. (A-F) Plot of the simulation of population growth (A, C, E) and linked average telomere length (B, D, F) as measured each 24 h, as schematized in Fig. 1A, with best-fit parameters of Table 1 and each day an initial number of cells *N*_*init*_= 1000, reaching a saturation number *N*_*sat*_ = *r*_*sat*_*N*_*init*_, with *r*_*sat*_ = 1000. HSL and relevant x-axis coordinates are indicated in red (A-B) Effect of initial global telomere length distribution translation towards longer or shorter average telomeres on replicative senescence at the scale of populations. (C-D) Effect of altering positively or negatively the left side of telomere length distribution. (E-F) Effect of increasing telomere-independent constant mortality rate (*p*_*accident*_). (G) Generation of senescence onset for simulated independent lineages as schematized in Fig. 1B ordered by lifespan using different constant mortality rates (*p*_*accident*_). (H) Median generation onset derived from (G) plotted as a function of increasing (*p*_*accident*_).

In order to provide context of numerous pathological conditions that demand increased cell turnover and renewal, we next evaluated the impact of a perturbation that is not directly linked to telomere processing by increasing the telomere-independent and constant mortality. We found a dramatic effect on senescence curves, with an anticipated HSL of ∼1 day at 20x increase in mortality, which corresponds to an increase from 0.43% estimated in wild-type cells, to 8.6% (Fig. 4E and S4E). Also, the apparent shortening rate in the population was found to increase significantly with increasing constant mortality (Fig. 4F and S4F). We concluded that the kinetics of replicative senescence and telomere shortening are remarkably influenced to co-morbidities.

Many mutants used to study telomere replication and replicative senescence, especially the ones involved in the repair of DNA damage, display varying degrees of slow growth or mortality, despite being considered viable, due to their involvement in telomere-independent aspects of genome stability. Yet, the fact that these mutations accelerate senescence has been interpreted as supporting their direct role in telomere maintenance. To determine whether our model could test the validity of the genetic interaction between telomerase inactivation and these mutants, we tested the deletion of *RAD51*, which encodes a recombinase that binds to short telomeres and is involved in the emergence of post-senescence survivors, and perhaps other mechanisms operating at short telomeres (*28, 37-39*). *RAD51* deletion has been described as accelerating senescence in populations to various extents (*37, 38, 40, 41*). However, it also increases the constant mortality to ∼5.4%, probably due to other functions in double-strand break repair or replication stress. To measure the contribution of the constant mortality caused by *RAD51* deletion in the observed accelerated senescence of *rad51*Δ in telomerase-inactivated conditions, we performed 1000 simulations of 11 individual cell lineages as if they were grown in the microfluidics device in the absence of telomerase and with a constant mortality rate of 5.4% and compared it to experimental data extracted from (*14*). We found that a large portion of the experimental curve fit within the confidence interval of simulations (Fig. 4G). We conclude that replicative senescence in the absence of *RAD51* is largely caused by telomerase-independent Rad51-dependent constant mortality. Because part of the curve lies outside the confidence interval, Rad51 may also specifically affect cells also when telomerase is absent. Hence, assessing synthetic lethality of *RAD51* deletion using replicative senescence as a readout may not be the most suitable approach for evaluating the genetic interaction between telomerase and *RAD51* gene.

## Discussion

In this study, we leveraged microfluidics-based data on individual lineages of dividing yeast cells with inactivated telomerase to build a comprehensive model of replicative senescence, and then applied it to simulate population assays commonly used to study replicative senescence. We combined information from previous studies in which we found that cell divisions are constrained by telomere shortening, and more specifically by the shortening of the shortest telomere in cells, and not simply by time or other telomere-related parameters, to fully account for replicative senescence heterogeneity (*16*). It is known that selection biases occur between lineage and population observations (*42-44*). Simulation enables drawing rigorous conclusions on population dynamics from lineage data observations. Our study contributes to this methodology, and allows us to decipher which phenomena and mechanisms prevail in populations, sometimes quite distinct and even counter-intuitive compared to what might be assumed at first sight from lineage data.

One of the most striking findings of this model lies in the contrasted route to senescence of *type A*, for which senescence occurs at an almost deterministic threshold of around 27 bp for the shortest telomere, versus *type B* cells, for which the probability to trigger senescence is telomere-length independent after experiencing a priming telomere-length-dependent non-terminal arrest. We thus suggest the existence of tight control mechanisms responsible for triggering senescence when a telomere becomes critically short in *type A* cells. Conversely, the mechanism signaling the terminal arrest in *type B* cells is likely distinct from the one operating in *type A* cells. *Type B* cells could die from indirect consequences of some processes initiated during the non-terminal arrests, either the prolonged DNA damage checkpoint activation and its genetic and epigenetic reprogramming, or the mutational burden associated with it.

In an experimental setting where one telomere is set shorter than the others and specifically sequenced, telomeres with lengths between 10 and 70 bp can be extracted from a senescing cell population, and even some chromosome ends lacking telomeric repeats can be detected (*15*). Recent experimental works measuring telomere length in telomerase-negative cells using long-read sequencing proposed an estimation of the threshold length for the critically short telomere around 70-75 bp (*28, 45*). When we take into account both *type A* and *B* cells in our model, we find an average length lower than 70-75 bp (Fig. 3H). However, we note that this average, and in particular the threshold length of 27 bp for *type A* cells which contributes to it, should not be taken as an exact value, but rather as an estimate which can depend on simplifying assumptions of the model, on the data used to fit and on uncertainties surrounding parameter estimation. It is important to note that for *type B* cells, we observed an accumulation of cells entering senescence at *ℓ*_*min,B*_ = 0, indicating that *in vivo, type B* cells might indeed be the carriers of telomere-free chromosome ends. We also refer to the works of (*45, 46*), which suggest that individual telomeres display differences in homeostasis length, possibly giving rise to different critical length thresholds. Furthermore, subtelomeric elements and heterochromatin status *in cis* might affect the critical length. Our simulations also indicate that the average shortest telomere length, as well as the average threshold length for senescence decreases over time because of competition and, as a consequence, the change of population structure, i.e. relative proportion of *type A* and *B* cells (Fig. 3A, C and Supp Fig. S3B).

When simulating population assays, we did not take into account post-senescence survivor emergence for two reasons: first, no microfluidics-based data was available to build and calibrate a model. Second, we wanted to focus on understanding to which extent replicative senescence on its own could explain experimental observations. Our study highlighted the presence of heterogeneity within the population, where cells of different generational ages and different histories co-exist, in particular *type B* cells, which have potentially experienced molecular and cellular events leading to genomic instability (*23*). Importantly, our observation that senescent cells are mainly *type A* between day 1 and 4, then after day 4, mainly *type B* (Fig. 3B, C) – an alternate dominance well-explained by their contrasted route to senescence – reinforce the idea that *type B* cells might be poised to generate post-senescence survivors. In this scenario, it is possible that the non-terminal arrests could correspond to attempts at DNA repair operating at the signaling telomere that could alter the length and structure of the shortest telomere itself, as suggested in (*47*).

Beyond the competition affecting population structure and dynamics, shifting perspective from individual lineages to populations reveals that, while the length of the initial shortest telomeres fully constrains senescence onset in lineages, the initial full distribution of telomere length, and not only the initial distribution of the shortest ones, is important for population growth dynamics (Fig. 4A and C). This is in accordance with the strong selection bias towards cells displaying the longest shortest telomeres in the initial population (Fig. 3E-G, S3A). We can also speculate that as time passes, senescence in a population is more often triggered by the telomeres which were initially ranked as second, third, fourth, etc. shortest telomere in the ancestor cell. A direct consequence of this dynamics is that the size and telomere length heterogeneity of the initial population prior to telomerase inactivation is critical to predict proliferation capacity. Conversely, if a specific critically short telomere is inherited in the same way in all cells, it is expected to have a significant impact on the whole progeny, in accordance with experimental results (*11, 15*). In the context of humans, for instance, in which telomerase activity is repressed in many somatic tissues, the genealogy of telomerase-positive stem cell compartments is thus expected to be key to determine how a given tissue proliferation and renewal capacity depends on the length of the shortest telomere rather than on the global distribution of telomere lengths.

Replicative senescence in the absence of telomerase has been extensively investigated in different genetic backgrounds and various conditions. Conditions where accelerated or delayed replicative senescence is observed, by comparing the kinetics of senescence at the population level, have often been interpreted as interfering with telomere biology (*25, 38, 48*). However, the fact that replicative senescence displays an intrinsic heterogeneity and a non-constant mortality makes it not trivial to draw such conclusions, as exemplified by the *RAD51* deletion effect on senescence, which we found to stem mostly from its intrinsic mortality and less from a specific requirement of Rad51 in the absence of telomerase (Fig. 4). In addition, relying solely on telomere length at specific time points or telomere shortening as predictors of senescence is inadequate, particularly when the initial telomere length is modified or mortality is influenced by factors other than telomerase inactivation.

Our mathematical model built based on a unicellular eukaryote paves the way for in-depth exploration of the intricate relationship between telomere length, shortening dynamics (*4*), and cell growth in different tissues in metazoans (as in (*49*)). It also provides a solid conceptual framework for the notion that accelerated telomere shortening and premature stem cell depletion may underlie various human diseases, particularly when compounded by co-morbidities that elevate cell mortality, such as the Duchenne muscular dystrophy (*50*). Our mathematical model might thus serve as a valuable approach for investigating the broader implications of telomere length dynamics in the context of human health.

## Materials and Methods

### Telomere shortening model

We use the model of (*16*), which consists in imposing that for each chromosome 1/ Only one of the two telomeres is shortened, the other conserves the parental length; 2/ It is shortened by the overhang, say h, assumed constant; 3/ The telomere shortened for one daughter is unchanged for the other daughter. Mathematically, at the *n*-th generation, we denote by 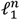 and 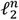 the random variables of the lengths of the two telomeric ends of a given chromosome at generation *n*. One and only one telomere is shortened by the overhang *h* with equiprobability, such that at the next generation for one of the two daughters we have 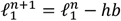 and 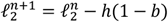, and for the other daughter it is the reverse: 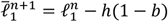 and 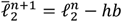, where *b*∼*Ber(1/2)* is a Bernoulli random variable coupling the two telomere lengths (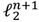 is shortened by *h* nucleotides if *b* is *0*, while 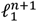 is left unchanged, and conversely if *b* equals *1*). Inside cells containing *2k* telomeres (*k=16*), we assume that telomeres of different chromosomes are independent (*51*), so that denoting by 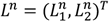 the matrix of size *2xk* of the lengths of the *2k* telomeres, we have similarly for the two daughters *L*^*n*^ and 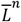

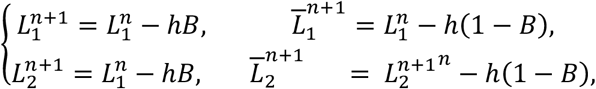

with *B* = (*B*_1_,…, *B*_*k*_) ∼ *Ber*(*k*, 1/2) is a random vector of *k* independent Bernoulli variables.

To model the population experiment (Fig. 1A), we keep the two daughters at each division, whereas for the microfluidic experiment (Fig. 1B), we pick up one of the two matrices 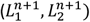 and 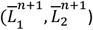 randomly uniformly.

### Initial distribution of telomere lengths

We assume that initially all telomere lengths are independent identically distributed (*i*.*i*.*d*.) according to a law 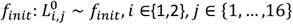. Given that generation *0* corresponds in our dataset to the inactivation of the telomerase, we depart from the distribution of telomere lengths in a telomerase-positive population (of the same yeast strain as the dataset) at equilibrium. We rely on the distribution of telomere lengths *f*_:_ of (*16*) (Fig. S1A) derived by adapting the numerical approach of (*11*) to this yeast strains. Given that the left-tail of the distribution has great influence on the lineages dynamics but is poorly characterized experimentally, we test small modifications of *f*_0_, namely translation (by a given length *ℓ*_*trans*_) and dilatations preserving the mode of the distribution, i.e. dilatations defined by dilating [*ℓ*_*inf*_, *ℓ*_*mode*_] (resp. [*ℓ*_*mode*_, *ℓ*_*sub*_]) to [*ℓ*_*inf*_+ *ℓ*_0_, *ℓ*_*mode*_] (resp. to [*ℓ*_*mode*_, *ℓ*_*mode*_ + *ℓ*_1_]) (see Supplementary Material for further detail). We optimize the values for *ℓ*_*trans*_, *ℓ*_0_ and *ℓ*_1_ together with the other parameters of the model, see Table 1 for the optimal values and Fig. S1A for the initial distribution *f*_*init*_.

### Laws of arrest

Once its telomere lengths determined by the telomere shortening model, the cell type is chosen according to the transition probability laws described by the tree diagram of Fig. 1E. The probability rate *p*_*accident*_ represents the chance to die accidentally and is constant, taken from (*23*). The rates *p*_*sen,A*_ and *p*_*sen,B*_are the probability to enter senescence respectively for *type A* and *type B* cells. They depend on the minimal telomere length *ℓ* = *min* (*L*^*n*^) through the law

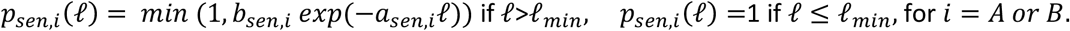

These laws are thus defined by five parameters (*a*_*sen,A*_, *b*_*sen,A*_, *a*_*sen,B*_, *b*_*sen,B*_, *ℓ*_*min*’_). Fig. 1F displays their best-fit values, which shows a remarkable fact: we can simplify the law for *type A* cells into an all-or-nothing law *p*_*sen,A*_=0 for *ℓ*>*ℓ*_*min*_ and *p*_*sen,A*_=1 for *ℓ* ≤ *ℓ*_*min*_, whereas *p*_*sen,B*_ (*ℓ*) = *b*_*sen,B*_*exp*(−*a*_*sen,B*_*ℓ*) is a very flat almost uniform law, so that finally only three parameters matter, namely *ℓ*_*min*_, *a*_*sen,B*_and *b*_*sen,B*_.

The rates for non-terminal arrests are similar: we define them through the same two-parameter law as for senescence: *p*_*nta*_(*ℓ*) = *min* (1, *b*_*nta*_ *exp*(−*a*_*nta*_*ℓ*)). Finally, when a cell experiences a sequence of arrests, (*24*) has successfully described the number of consecutive abnormally long cycles, either terminal or not, by a geometrical law. This corresponds to a constant probability to exit this sequence either by repairing or adapting (*p*_*repair*_) after a non-terminal arrest, or at the opposite by dying (*p*_*death*_)after a senescent cycle.

### Laws of cell cycle duration times

Whereas cell cycle durations are not fundamental to simulate the microfluidic experiments, the definition of laws for the duration of cell cycles is crucial to model the population experiment. Based on experimental measurements, and on a threshold *D* above which a cell cycle duration is considered abnormally long, we estimated the laws in the four distinct regimes: normal cell cycle durations for *type A* and *type B* cells, non-terminal arrest cycle durations and senescent cycle durations (Fig. 1G). According to the cell type and cell cycle type, the cycle duration is then picked up at random following the corresponding law.

### Parameters values

#### Parameters from the literature

The shortening length is taken as the overhang length *h* = 7 bp, see (*20*). We define the threshold for abnormally long cell cycle durations *D* = 180 min, following the thorough sensitivity analysis carried out in (*24*). From the same reference, we fixed *p*_*death*_ = 0.58 and*p*_*repair*_ = 0.65. From (*23*) we get *p*_*accident*_ = 4.3. 10^−3^.

#### Parameters from direct experimental measurements

In our experiments we have measured *r*_*sat*_ = 720. We also pick up at random the cell cycle durations fron the experimental distributions, classified into the four categories displayed in Fig. 1G.

### Determination of Ninit

The experimental value for *N*_*init*_ is around 10^5^, which is out of reach for simulations. We thus compare the behavior of populations originating from different initial number of cells *N*_*init*_. In order to accurately estimate these behaviors (i.e. to have an empirical behavior close to the statistical one) we simulated 25 times the evolution of a population with a certain fixed initial size *N*_*init*_. The full sensitivity analysis is detailed in Supplementary Material, see also Fig S2. We first noticed that the less cell initially present, the more variability between simulations, which supports the idea that the variability in the distribution of initial telomere lengths is an important source of heterogeneity in senescence (*11, 15*). Even though extreme behaviors are sensitive to *N*_*init*_ unlike average behaviors, the decrease in variability stabilizes around *N*_*init*_ = 200, and then corresponds to the variability intrinsic to the stochastic evolution.

## Acknowledgements

We thank Claus Azzalin, Miguel Godinho Ferreira, and Helen Pickersgill from Life Science Editors for critical reading of the manuscript.

## Author contributions

Conceptualization, Funding acquisition, Supervision: MD, MTT, ZX; Methodology: AR, MD, ZX; Investigation & Writing: AR, MD, MTT, ZX.

## Data and materials availability

Codes have been deposited at https://github.com/anais-rat/telomeres

## Supplementary Materials

Figs. S1 to S4

Extended Materials and Methods

