## Supplementary Materials for "Individual cell fate and population dynamics revealed by a mathematical model linking telomere length and replicative senescence"

This pdf includes:

Supplementary Figures S1-S4

Extended Materials and Methods

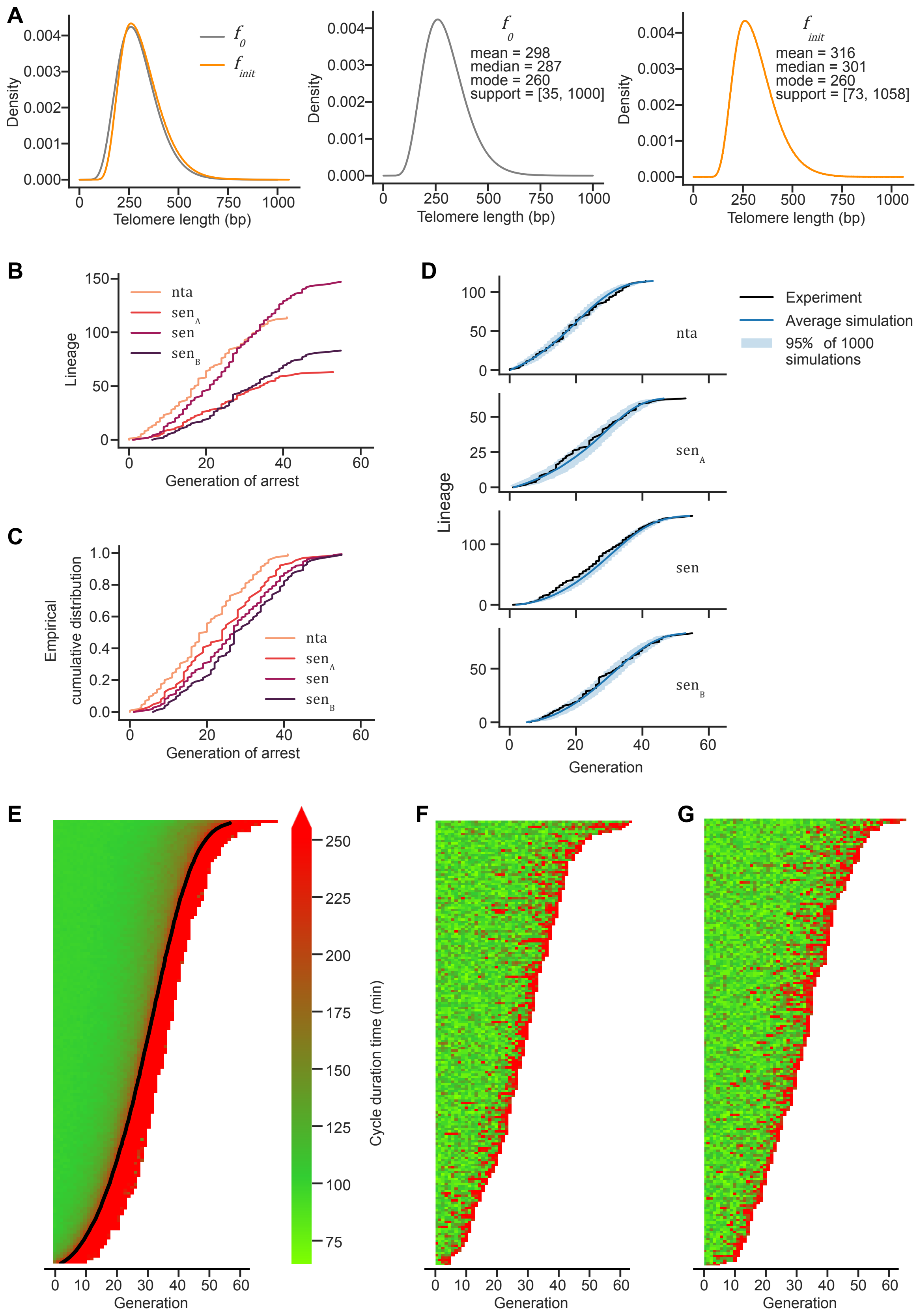

A
Density
Telomere length (bp)
$f_0$
$f_{init}$
mean = 298
median = 287
mode = 260
support = [35, 1000]
mean = 316
median = 301
mode = 260
support = [73, 1058]
B
Lineage
Generation of arrest
nta
sen$_A$
sen
sen$_B$
C
Empirical cumulative distribution
Generation of arrest
D
Experiment
Average simulation
95% of 1000 simulations
Lineage
Generation
E
Generation
Cycle duration time (min)
F
Generation
G
Generation

#### *Figure S1: Model description details*

(A) Initial telomere lengths distributions. $f_0$ has been computed in (*16*) by adapting the approach of (*11*) to a yeast strain expressing telomerase RNA subunit under a conditional promoter as used in microfluidics experiments (*14, 52*). $f_{init}$ corresponds to the slight modification of $f_0$ described in Methods with parameters of Table 1. (B) Experimental generation of arrest classified by lineage *type* and type of arrest (non-terminal or senescent arrest, as extracted from Fig. 1C. *nta*: generation of the first non-terminal arrest; $sen_A$, $sen_B$: generation of first senescent arrest for *type A, B* cells; *sen*: generation of first senescent arrest for all *types*. (C) same as in (B) normalized to obtain empirical cumulative distribution. (D) Comparison between 1000 random simulations of the best-fit model and experimental data displayed in (B). (E) Average over 1000 random simulations of the microfluidics experiment using the best-fit model (parameters in Table 1). (F, G) Simulated data of one microfluidics experiment with the best-fit model, as in Fig. 1H.

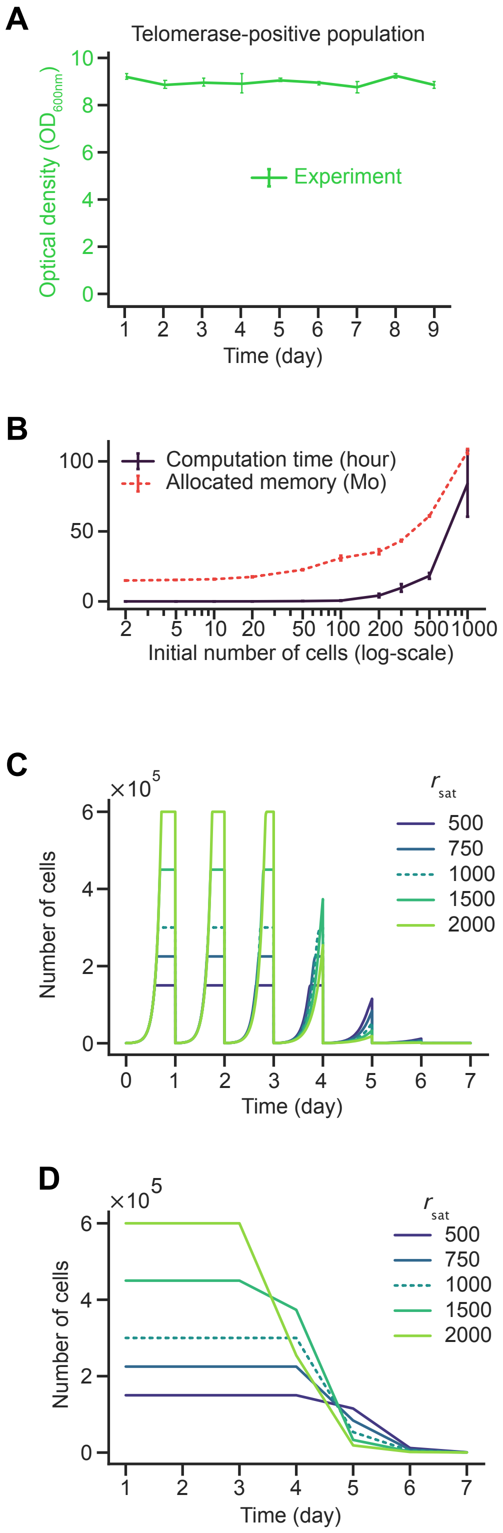


*Figure S2: Study to establish the parameters for the population simulation*

(A) Experimental data of cell growth as in Fig. 2B in telomerase-positive conditions to determine the ratio $r_{sat}$ to generate Fig. 2B. (B) Computation time and allocated memory with respect to $N_{init}$. (C) Saturation time since last dilution with respect to $N_{init}$ in days 1 to 4. (C-D) Population growth as a function of time (C) or as measured every 24h (D) for indicated $r_{sat}$ values.

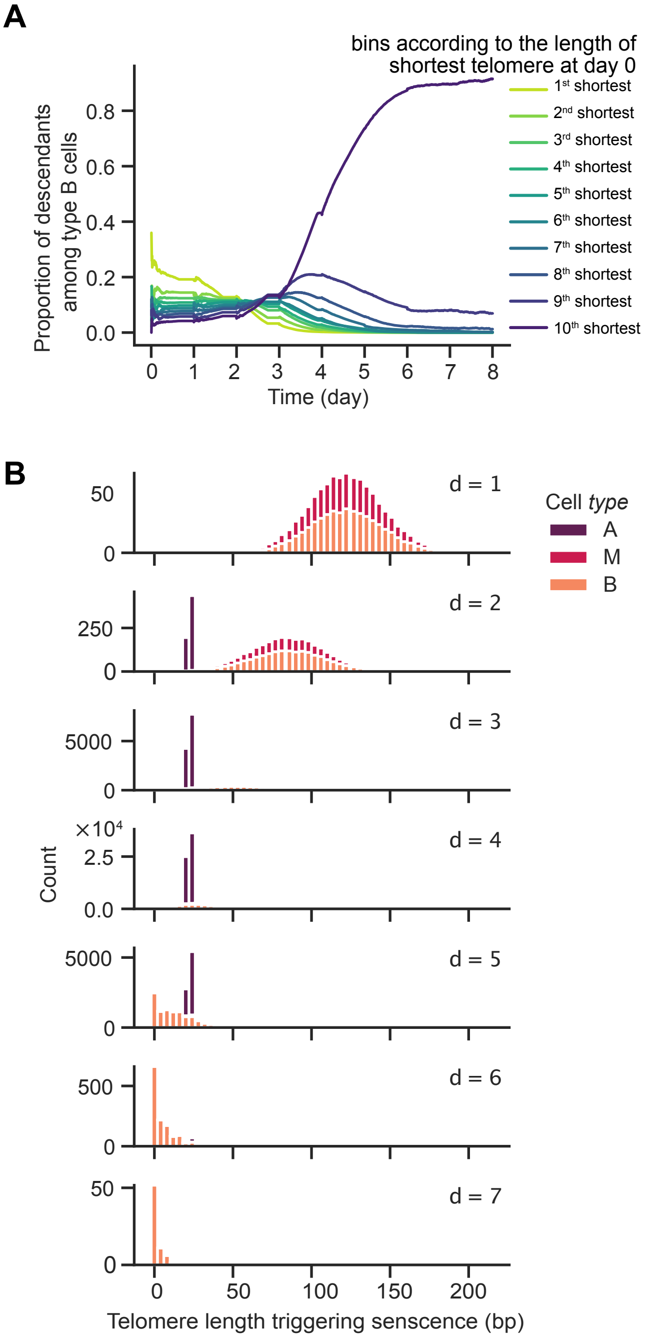


*Figure S3: Additional hidden parameters, experimentally inaccessible, extracted from simulations of telomerase-negative cell cultures.*

(A) Same as (Fig. 3E) for *type B* cells only. (B) Distribution of the length of the shortest telomere in senescent cells for indicated cell *types* for each day.

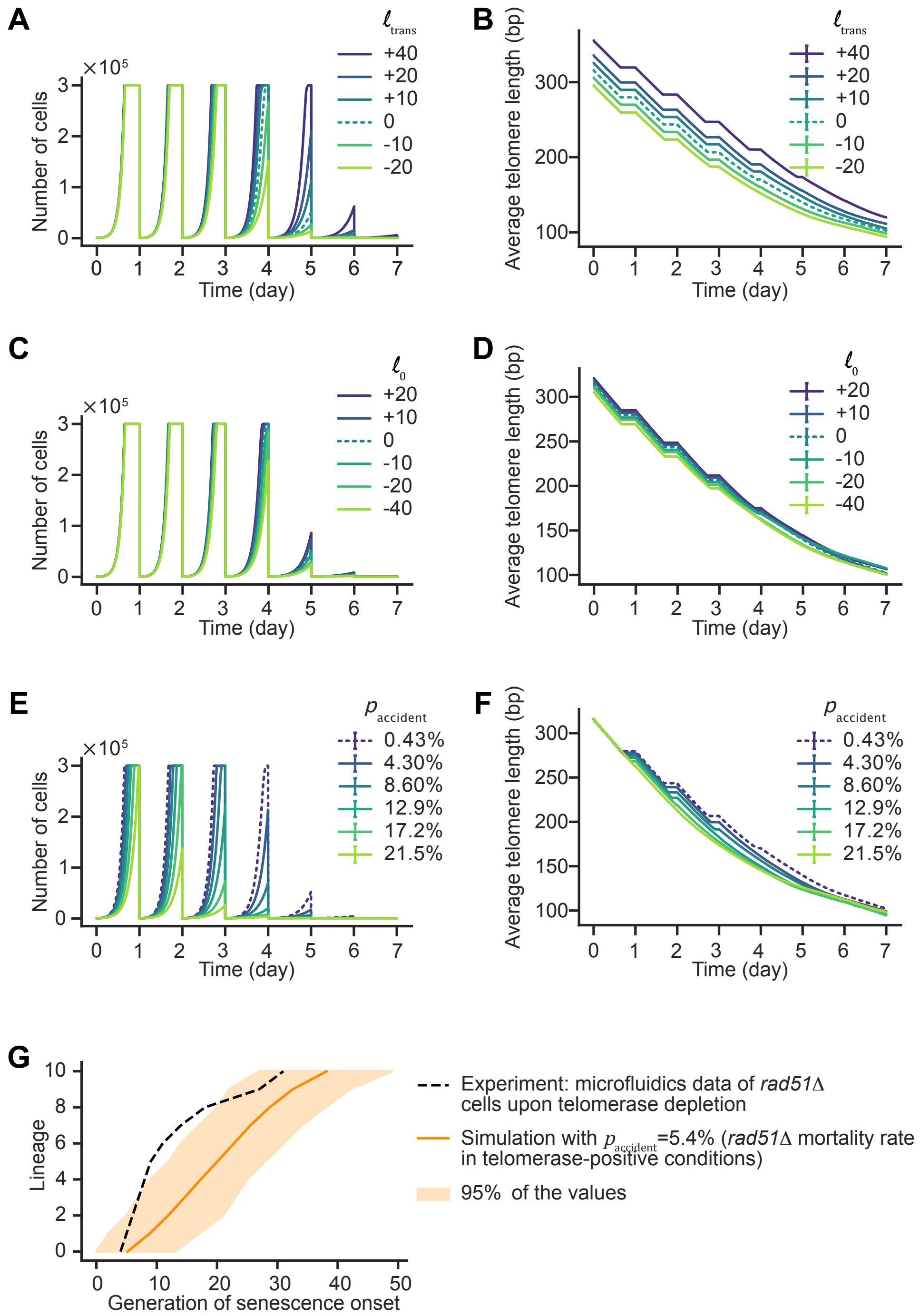


*Figure S4: Effects of altering telomere length prior to telomerase inactivation or the constant mortality rate on replicative senescence.*

(A-F) Same as Fig. 4A-F but plots correspond to all values simulated with time (and not values reached each day) (G) Lifespan of individual cell lineages of *rad51Δ* cells upon telomerase inactivation, as assessed by fmicrofluidics (from (*14*)) and 1000 simulations of 11 individual cell lineages, corresponding the number of experimental lineages, using indicated $p_{\text{accident}}$.

### Extended Materials and Methods on mathematical modeling, numerical simulation and parameter estimation of the article: *Individual cell fate and population dynamics revealed by a mathematical model linking telomere length and replicative senescence*

Anaïs Rat Marie Doumic Teresa Teixeira Zhou Xu

November 20, 2023

In these extended Materials and Methods, we provide a fully detailed description on the mathematical modeling, the numerical simulations and the parameter estimation methodology we developed. For the sake of clarity and completeness, we have included supplementary figures which are not cited in the Main Text. When referring to "Fig. 1", we thus mean "Fig. 1 of this extended Materials and Methods"; to refer to Fig. 1 of the Main Text, we refer to "Main Fig. 1".

The codes are available at `https://github.com/anais-rat/telomeres`.

#### 1 Initial distribution of telomere lengths

**Departure point: a telomerase-positive telomere shortening/elongating model** Given that generation 0 corresponds in our dataset to the inactivation of the telomerase, the inital distribution of telomere lengths corresponds to the distribution of telomere lengths in a telomerase-positive population of the same yeast strain as the dataset. It is natural to assume that in such a population, a steady distribution of telomere lengths has been reached, and is the same for any telomere of the cells. For this reason, we assume that initially all telomere lengths are independent, identically distributed (i.i.d.) according to the same law $f_{init}$:

$$L^0_{i,j} \sim f_{init}, \qquad i \in \{1,2\}, \qquad j \in \{1,\dots,16\},$$

where $L^0_{i,j}$ is the random variable corresponding to the length of the $i$th telomere of the $j$th chromosome at generation 0.

To determine $f_{init}$, we thus rely on the distribution of telomere lengths $f_0$ of Bourgeron et al. [3] (Fig. S1A) derived by adapting the numerical approach of [7] to the yeast strain that interests us. Their distribution approaches the stationary distribution of a certain Markov chain, whose parameters are fitted on experimental data, which describes telomere length shortening and elongation in the presence of telomerase.

**Sensitivity analysis on the initial telomere length distribution** Due to the fact that very few experimental measurements of the telomere length distribution $f_0$ are available, so that it has been estimated indirectly from the mathematical shortening/elongation model described above and the experimental measurement of its statistical mode, we allowed $f_{init}$ to deviate slightly from $f_0$, namely:

- translations by $\ell_{trans}$,
- dilations at both sides of the mode which preserve the (experimentally measured) mode:
$$\begin{cases} \inf\big(\mathrm{supp}(f_{init})\big) = \ell_{inf} + \ell_0 \\ \sup\big(\mathrm{supp}(f_{init})\big) = \ell_{sup} + \ell_1 \\ \mathrm{mod}(f_{init}) = \ell_{mode} \end{cases}$$
to preserve the mode $\ell_{mode} \coloneqq 260(16)\,\mathrm{bp}$ [8] of the experimental distribution and bring $[\ell_{inf}, \ell_{mode}]$ on $[\ell_{inf} + \ell_0, \ell_{mode}]$ and $[\ell_{mode}, \ell_{sup}]$ on $[\ell_{mode}, \ell_{sup} + \ell_1]$.

Finally, the transformation of $f_0$ can be written as

$$f_{init}\big(\cdot; \ell_{trans}, \ell_0, \ell_1\big) = C \times \begin{cases} f_0\big((\ell - \ell_{inf} - \ell_0)\alpha_0 + \ell_{inf} - \ell_{trans}\big) & \ell \in \big[\ell_{inf} + \ell_0, \ell_{mode}\big]\,, \\ f_0\big((\ell - \ell_{mode})\alpha_1 + \ell_{mode} - \ell_{trans}\big) & \ell \in \big[\ell_{mode}, \ell_{sup} + \ell_1\big]\,, \end{cases} \tag{1.1}$$

with $C = C(\ell_0, \ell_1)$ a normalization constant, and

$$\alpha_0 \coloneqq \frac{\ell_{mode} - \ell_{inf}}{\ell_{mode} - \ell_{inf} - \ell_0}, \qquad \alpha_1 \coloneqq \frac{\ell_{sup} - \ell_{mode}}{\ell_{sup} + \ell_1 - \ell_{mode}} \tag{1.2}$$

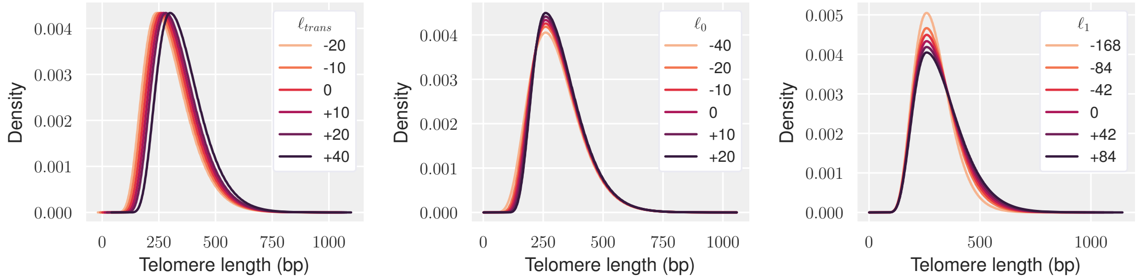


Figure 1: Transformations of the distribution of telomere lengths $f_0$: translations *(left)*, dilatation of the left tail *(middle)* and the right tail *(right)*.

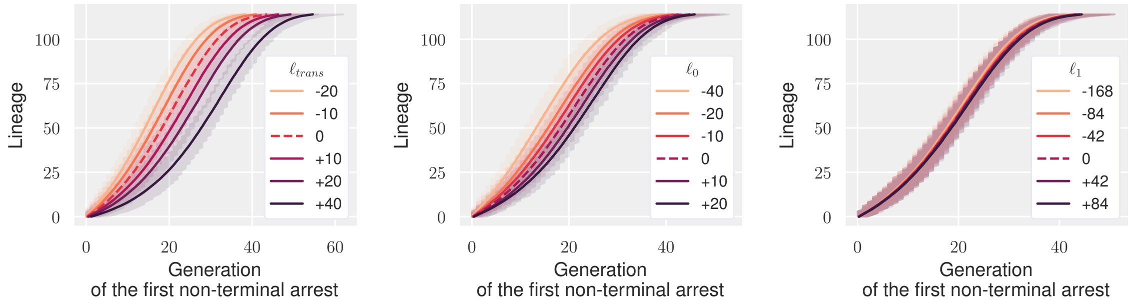


Figure 2: Generation of the onset of the first non-terminal arrest with respect to the transformations of $f_0$ displayed in Fig. 1: translations *(left)*, dilatation of the left tail *(middle)* and the right tail *(right)*. Average and $5^{\text{th}}$ and $95^{\text{th}}$ percentiles on $k = 1000$ simulations.

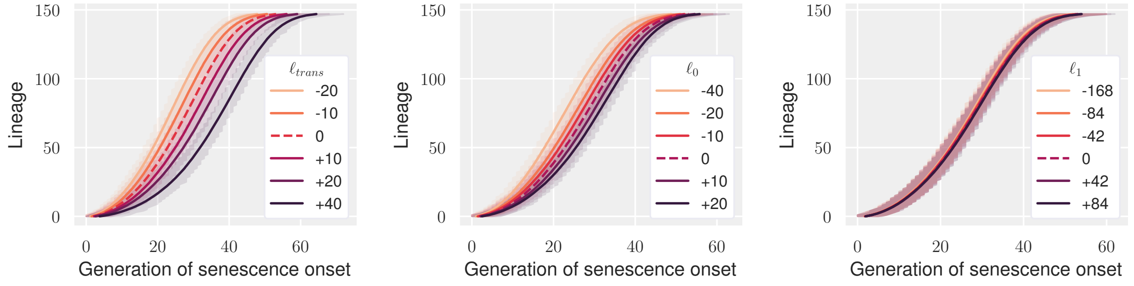


Figure 3: Generation of the onset of senescence with respect to the transformations of $f_0$ displayed in Fig. 1: translations *(left)*, dilatation of the left tail *(middle)* and the right tail *(right)*. Average and $5^{\text{th}}$ and $95^{\text{th}}$ percentiles on $k = 1000$ simulations.

In Fig. 1, modifications on $f_{init}$ are displayed, and in Fig. 2 and 3 their corresponding influence on the onset of non-terminal arrests (Fig. 2) and senescence (Fig. 3). The left tail of the distribution appears to have a great influence on the dynamics of microfluidic data - contrarily to its influence on the population experiment, see Main Text and Main Fig. 4D. We also see that the right tail dilatation has only little influence.

These modifications are then fitted together with the laws of arrest on microfluidics data; the distribution $f_{init}$ retrieved is plotted on Fig. S1A.

#### 2 Determination of $N_{init}$

In the population experiment, the initial concentration and the concentration of dilution are the same. Thus, if our simulations are initiated with $N_{init}$ cells, dilution consists in selecting $N_{init}$ cells *uniformly* among the population reached before dilution.

As for saturation, we consider that when the number of individuals becomes too high, all individuals stop dividing/evolving. To reach saturation we fix a population size $N_{sat}$ at which the population saturates. Since we cannot reasonably reach the experimental concentration *in silico* we rather fix the ratio $r_{sat}$ such that $N_{sat} = r_{sat}N_{init}$. In the experimental dataset $r_{sat} \approx 720$ whereas in other experiments it may be much larger.

**Representativity analysis – sensitivity to $\mathbf{N_{init}}$.** We compare the behavior of populations originating from *different initial number of cells $N_{init}$*. In order to accurately estimate these behaviors (i.e. to have an empirical behavior close to the statistical one) we simulated $k = 25$ times the evolution of a population with a certain fixed initial size $N_{init}$.

**Variability decreases with $N_{init}$ up to a certain point.** The first, expected, observation is that *the less cells initially present, the more variability between simulations*, see Figs. 4, 5, 7,9, 10 and Table 1. This supports the idea that the variability in the distribution of initial telomere lengths is an important source of heterogeneity in senescence [1, 7]. The decrease in variability however seems to stabilize around $N_{init} = 200$, at a level which corresponds to the intrinsic variability, linked to the stochastic nature of the dynamics.

**Extreme behaviors are sensitive to $N_{init}$ unlike average behaviors.** Starting initially with more cells, we are more likely to hit the tails of the distribution of initial telomere lengths and to have chromosome shortening mainly on the same extremity, resulting in a fast shortening telomere and a slow shortening one. Extremum quantities are thus sensitive to $N_{init}$. This can be observed for example looking at the evolution of the minimum telomere length in the population plotted on Fig. 4. The same reasoning applies to the extinction time of the population, which is the maximal lineage lifetime in the population:

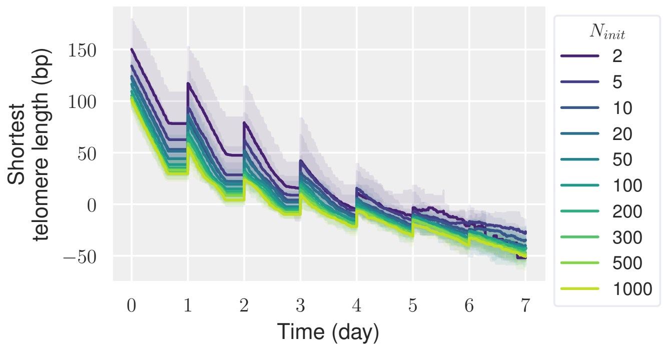


Figure 4: Time evolution of the minimal telomere length in the population with respect to the initial number of cells $N_{init}$. Average and $5^{th}$ and $95^{th}$ percentiles on $k = 25$ simulations. (The plateau sections correspond to saturation, during which the population stops evolving.) Although decreasing with $N_{init}$, the shortest-telomere length seems to converge.

it keeps increasing with the initial number of cells, and so is the senescing time, see Fig. 5.

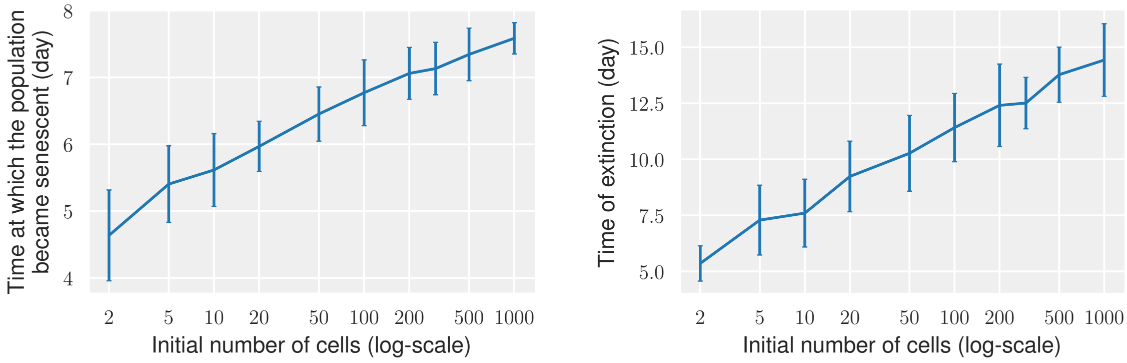


Figure 5: Graphs of the times of senescence *(left)* and extinction *(right)* of simulated populations with respect to the initial number of cells $N_{init}$. Average and standard deviation on $k = 25$ simulations. Although senescence and extinction times keep increasing with $N_{init}$, the log-scale indicates that the increase tends to zero.

Since the length of the shortest telomere is a major determinant of the onset of senescence, Fig. 5 and 6 can be explained looking at the distributions $f^n_{min}$ and $f^n_{min-max}$ of the minimum and maximum, respectively, of the shortest telomere in a population of $n$ cells (each cell possessing 32 telomeres). As shown on Fig. 6, they get translated and narrowed for large $n$. To access $f^n_{min}$ and $f^n_{min-max}$ we compute their cumulative distribution functions, benefiting from the independence of telomere lengths at generation 0:

$$\begin{aligned} F^n_{min}(\ell) &= \mathbb{P}\Big(\min_{1\le i\le n}\Big(\min_{1\le j\le 32} L_{i,j}\Big) \le \ell\Big) = 1 - \mathbb{P}\Big(\bigcap_{\substack{1\le i\le n\\1\le j\le 32}} L_{i,j} > \ell\Big) \\ &= 1 - \big(1 - F_{init}(\ell)\big)^{32n}, \end{aligned}$$

where $F_{init}$ denotes the cumulative distribution function of $f_{init}$, and

$$\begin{aligned} F^n_{min-max}(\ell) &= \mathbb{P}\Big(\max_{1\le i\le n}\Big(\min_{1\le j\le 32} L_{i,j}\Big) \le \ell\Big) = \mathbb{P}\Big(\bigcap_{1\le i\le n}\Big(\min_{1\le j\le 32} L_{i,j}\Big) \le \ell\Big) \\ &= F^1_{min}(\ell)^n = \Big(1 - \big(1 - F_{init}(\ell)\big)^{32}\Big)^n, \end{aligned}$$

from which we deduce

$$\begin{cases} f^n_{min}(\ell) = F^n_{min}(\ell) - F^n_{min}(\ell-1) = \big(1 - F_{init}(\ell-1)\big)^{32n} - \big(1 - F_{init}(\ell)\big)^{32n}, \\ f^n_{min-max}(\ell) = \Big(1 - \big(1 - F_{init}(\ell)\big)^{32}\Big)^n - \Big(1 - \big(1 - F_{init}(\ell-1)\big)^{32}\Big)^n. \end{cases} \tag{2.1}$$

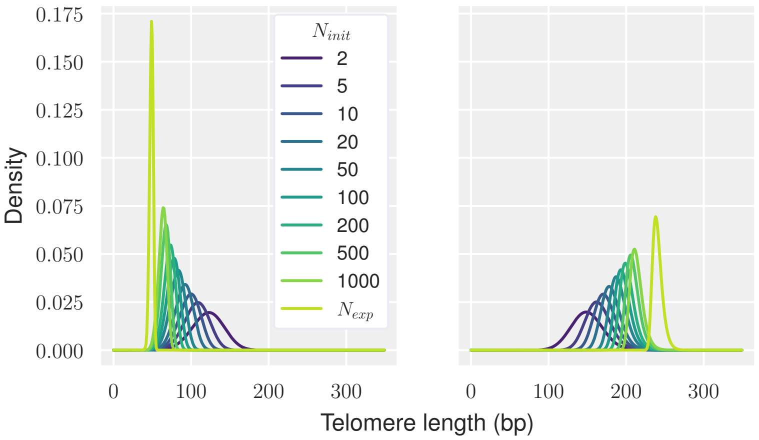


Figure 6: Distribution of the minimal *(left)* and maximal *(right)* shortest telomere length in a population of $N_{init}$ cells.

The evolution of average indicators are less sensitive to $N_{init}$ than the time evolution of extrema, as for example

- the average generation (Fig. 7-*right*),
- the average over all cells of the average telomere length, or even of the shortest telomere length (Fig. 7-*left*).

These quantities vary less with respect to $n$.

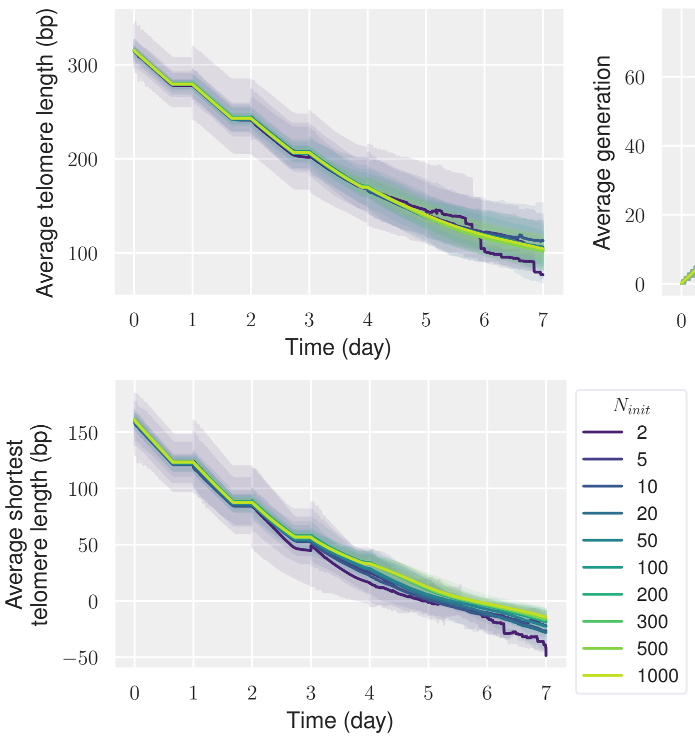


Figure 7: Time evolution of three population average quantities with respect to the initial number of cells $N_{init}$: the generation *(right)*, cells' average *(top left)* and shortest *(bottom left)* telomere length. Average (and $5^{\text{th}}$ and $95^{\text{th}}$ percentiles at the left) on $k = 25$ simulations.

**Saturation bias.** Likewise, the time when the saturation limit is reached is a time indicator less sensitive to the extreme behaviors, and thus more relevant (in the Main Text we also compare, during the sensitivity analysis, the time when half of the saturation limit is reached (HSL)). This saturation time is directly linked to the doubling-time of the population. On Fig. 8 we see that it stabilizes in average for $N_{init}$ greater than 50 and in standard deviation around $N_{init} = 200$, in accordance with Table 1.

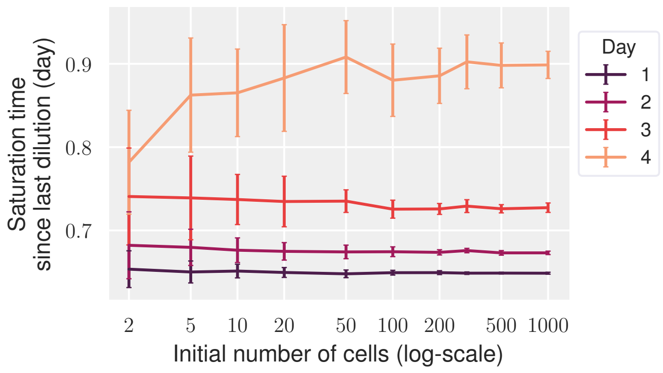


Figure 8: Graph of the times of saturation (on each day of saturation) of simulated populations with respect to the initial number of cells $N_{init}$. Average and standard deviation among the simulations that have reached saturation among $k = 25$ simulations.

| $\mathbf{N_{init}}$ | **2** | **5** | **10** | **20** | **50** | **100** | **300** | **500** | **1000** |
|---|---|---|---|---|---|---|---|---|---|
| **Day 1** | 1 | 1 | 1 | 1 | 1 | 1 | 1 | 1 | 1 |
| **Day 2** | 1 | 1 | 1 | 1 | 1 | 1 | 1 | 1 | 1 |
| **Day 3** | 0.84 | 0.96 | 1 | 1 | 1 | 1 | 1 | 1 | 1 |
| **Day 4** | 0.24 | 0.44 | 0.4 | 0.48 | 0.72 | 0.96 | 1 | 1 | 1 |

Table 1: Table of the proportion of simulations (among $k = 25$ simulations) that have saturated per day with respect to the initial number of cells $N_{init}$.

Having a closer look, we notice that not only small populations die and senesce earlier than large ones (Fig. 5), but they also saturate slightly later on the first days of saturation and they saturate "less" on the third and fourth day than large populations (Fig. 8 and 9, Table 1). In addition, the few small populations that have saturated on the fourth day have saturated earlier than large populations. Let us explain these observations. Given

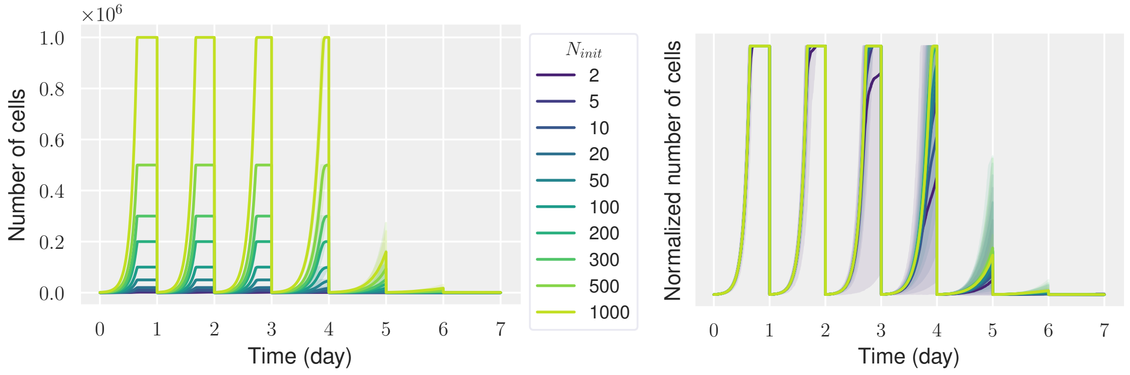


Figure 9: Time evolution of the number of cells with respect to the initial number of cells $N_{init}$. The number of cells through time is the effective one at the *(left)* and it is renormalized (by $\frac{N_{init}}{500}$) at the *(right)*. Average and 5$^{\text{th}}$ and 95$^{\text{th}}$ percentiles on $k = 25$ simulations.

our model, the evolution of cells is independent from each other. Under growth, division and death processes, any given subpopulation is therefore evolving independently from the rest of the population. However, cells become coupled through the total number of cells as soon as we add saturation at threshold $N_{sat}$ (all the cells stop evolving as soon as the total number of cells reaches $N_{sat}$). A population with a small pool of fast/slow dividing lineages will then reach saturation –and thus stop evolving– earlier/later than an homogeneous population, with the consequence of decreasing/increasing in the other part of the population the average generation at the next dilution. This explains why most of the small populations, by saturating late on the two first days, avoid saturation on the following days. Because cells have stopped evolving later, they are "closer to death" in accordance with early extinction time, and high proportion of senescent cells compared with large populations (see Fig. 10-*right*). On the contrary, the few small populations that have saturated on the 3$^{\text{rd}}$ and 4$^{\text{th}}$ days, the same that managed to reach day 7, are made of lineages that have undergone arrest: the proportion of *type B* cell on the last day (an average only among non-extincted populations) is particularly high (Fig. 10-*left*) and the average generation particularly low (Fig. 7-*right*).

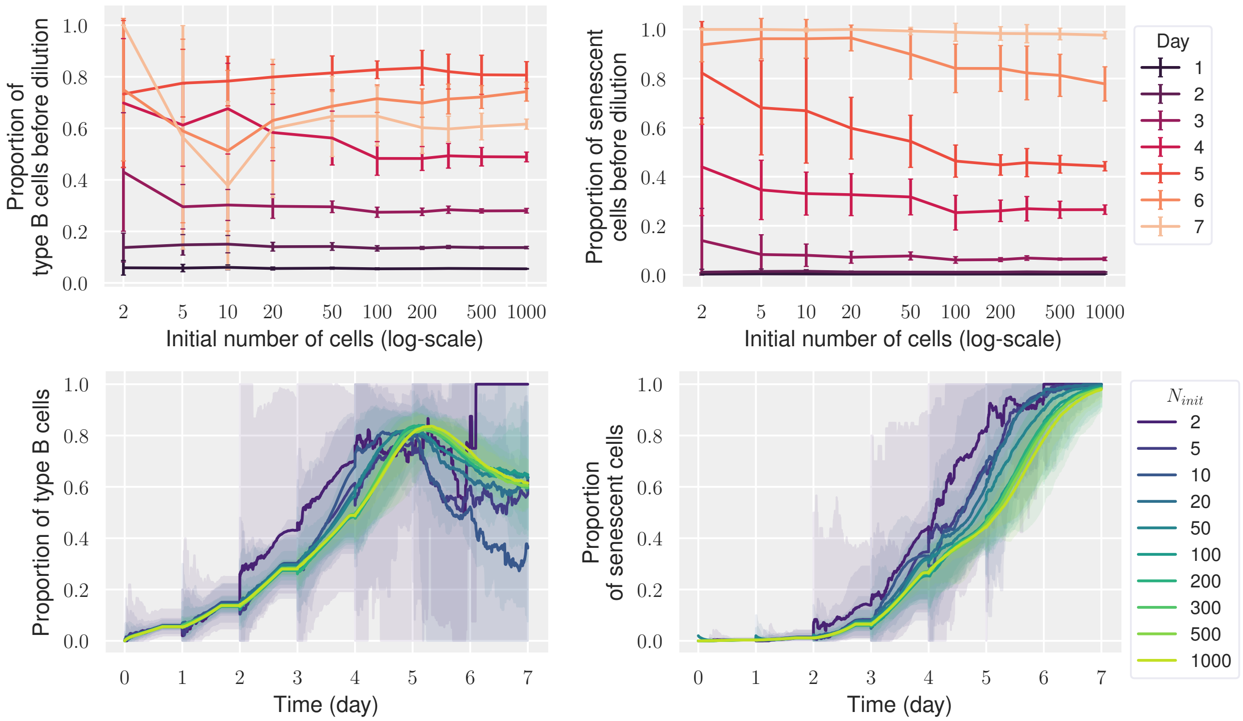


Figure 10: Proportion of *type B* cells *(left)* and senescent cells *(right)* in the population before each dilution *(top)* or at each time of the experiment *(bottom)* with respect to the initial number of cells $N_{init}$. Average and standard deviation *(top)* or 5$^{\text{th}}$ and 95$^{\text{th}}$ percentile *(bottom)* on $k = 25$ simulations. At each time, these statistics are taken only on those of the $k$ simulations that are not extincted, which bias the graphs of small populations at large times, made as an average on very few simulations.

**Dilution bias.** Such a saturation bias is accentuated by dilution since a small pool of fast/slow dividing lineages will be over/under represented right before dilution. Therefore the rest of the population will have less/more chance to be selected by dilution than it had been in an homogeneous population.

The dilution bias is significant only for the (small) populations that present a lot of variability in the saturation time. As already noticed, starting from around $N_{init} = 200$ the initial population is large enough to have only small fluctuations of the saturation time.

### 3 Implementation

The implementation of the model was coded in the Python programming language (Python Software Foundation, https://www.python.org/), with 3.8 Python version. The whole project is available at https://github.com/anais-rat/telomeres.

### 4 Inference of the parameters with microfluidics data

In the present section we present how we fit the parameters of the laws $p_{nta}$, $p_{sen_A}$ and $p_{sen_B}$ that characterize the onset of non-terminal and terminal arrests, respectively, enabling small transformations of the distribution $f_0$ of initial telomere lengths. We rely on microfluidic data and the simulation of lineages.

#### 4.1 Description of the estimation method

**Experimental data to fit.** To fit our models, we consider the data plotted on Fig. 11-*left*, namely:

1. $G_{sen}$ the generations at which senescence occurred, in the $n_{sen} = 148$ *experimentally senescent*, *i.e.* terminated by a long cycle and death, lineages of the dataset.
2. $G_{sen_A}$ the subset of the generations $G_{sen}$ of senescence made only of the $n_{sen_A} = 64$ (dead senescent) lineages classified as *experimental type A*. These are the lineages composed uniquely of normal cycles and experimental senescence.
3. $G_{sen_B}$ similarly with the remaining $n_{sen_B} = 84$ lineages of *experimental type B*. These are linages presenting at least one arrest followed by a normal cycle.
4. $G_{nta}$ the generations at which occurred a first non-terminal arrest, generated from a total of $n_{nta} = 115$ experimental *type B* lineages.
5. The proportion of the experimentally senescent lineages classified as *type B*:

$$r_{B\%} = 100 \times \frac{n_{sen_B}}{n_{sen}} \approx 57\,\%.$$

*Notations. In the following $G_{sen}(j)$ denotes the generation of senescence onset in the $j$th senescent lineage of the dataset (when ordering by increasing generation of senescence) and similarly with the types of arrest ("nta", "sen$_A$" and "sen$_B$"). Therefore the maps plotted on Fig. 11-left correspond to $G^{-1}$, but we abusively call them G-graphs (or* generation graphs*). Wee keep this choice by sake of coherence with the previous articles [3, 4].*

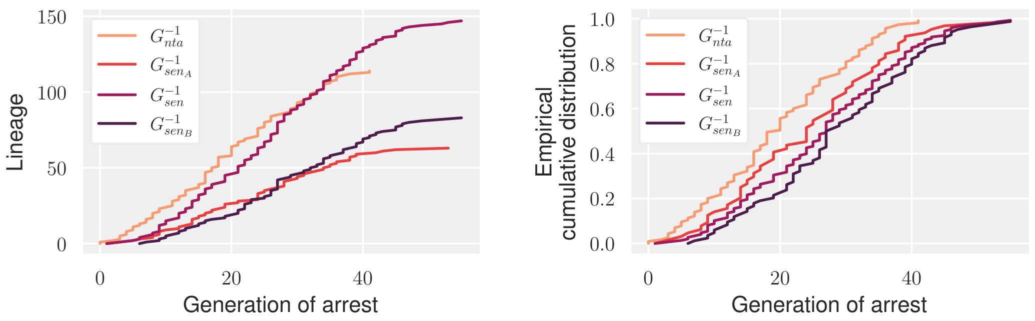


Figure 11: Representation of the experimental generations ($x$-axis) at the onset of a first non-terminal (nta) or terminal (sen) arrest, among different lineages (indexed on the y-axis by increasing generation of arrest) of same characteristics (classified as $A$ or $B$ *type* and/or senescent) *(left)*. They are obtained from the lineage dataset (see Main Fig. 1C of the Main Text) after classifying every cell cycle as either long or normal with threshold $D = 180\,\text{min}$. Every lineage is classified depending on the location of its long cycles and we extract the generation(s) at which it got arrested. For every generation graph, renormalizing the y-axis yields the empirical cumulative distribution function of the experimental law of the associated generation of first arrest *(right)*.

**Simulated "generation graphs".** To these experimental $G$-graphs correspond theoretical graphs given by our model. We approach these theoretical graphs with Monte Carlo estimators $\hat{G}^{(k)}$, realized as an average on $k$ independent simulated graphs $(\hat{G}^1, \dots, \hat{G}^k)$:

$$\hat{G}_i^{(k)} : j \mapsto \frac{1}{k} \sum_{s=1}^{k} \hat{G}_i^s(j), \qquad i \in \{sen,\ sen_A,\ sen_B,\ nta\},\ j \in \{1, \dots, n_i\}. \tag{4.1}$$

Each of the $k$ simulated graphs presents the same characteristics as the associated experimental graph: it is made of the same number of generations, similarly ordered and with same characteristics (i.e. describing the onset of a specific type of arrest in a specific type of lineage).

**Dependencies between simulated graph and parameters.** A delicate point is that simulated data depend on both the law for non-terminal arrests and senescence, and slightly on $p_{repair}$. In our model, we test at each division whether or not a newborn cell enters senescence (if its mother is not senescent) or enters or exits a non-terminal arrest. In particular, to enter a first non-terminal arrest at generation $g$, a lineage must have reached generation $g$ without having undergone any arrest. Thus, $\hat{G}_{nta}$ depends not only on the probability to experience a first non-terminal arrest but also on the probability to enter senescence.

Likewise, for a fixed law of senescence onset, if for example $p_{repair}$ is small and $p_{nta}$ is such that the entry in senescence is much more likely than the entry in a sequence of non-terminal arrest(s), very few lineages will have time to enter and exit a non-terminal sequence of arrests before entering senescence, with great impact on the proportion of experimental *type B* lineages and the generation graphs. This is mainly due to the fact that, in order to generate the graphs of generations approaching the experimental graphs, the generations of arrest and thus the type of lineages are considered as they would have been perceived experimentally rather than as they are in the model (other said, *type M* lineages are classified as *type A* for comparison with experimental data, see Main Text). For our best-fitted parameters, around 16 % of senescent lineages are misclassified *type A* which is not negligible.

**Parameters to estimate.** Even though the parameters of $p_{nta}$ were fitted by Martin et al. [4] with the same dataset, they were fitted with a fixed law of senescence entry, consisting in a deterministic threshold at 0 bp. Because of the dependency of the simulated graphs on both laws, we need to estimate $p_{nta}$ again, simultaneously with $p_{sen_A}$ and $p_{sen_B}$.

Still, we do not re-estimate $p_{repair}$, the probability to exit a non-terminal arrest given that senescence is not triggered. It was fitted by Martin et al. [4] with the distribution of the number of consecutive non-terminal arrests among all the sequences of non-terminal arrests. Modifying their laws of arrest in a way that fits the experimental curves does not modify significantly the distribution of consecutive long cycles (and thus $p_{repair}$). We verified this a posteriori with the best-fit laws of arrest.

**Method.** We used the evolutionary algorithm CMA-ES in its 3.2.2. python version available on GitHub at `https://github.com/CMA-ES/pycma`. Based on the principles of biological evolution, CMA-ES explores in a stochastic and evolutionary-driven way a certain domain $\mathcal{D}$ of parameters in order to estimate the point(s) of the domain that minimizes a given cost function $E$ [2]. For more details on the algorithm and on the CMA-ES parameters that we have adjusted see Table 2 or directly the GitHub link.

The cost function is defined as a weighted sum of the errors, in $\ell^2$-norm, between the four experimental $G$-graphs[1] and their estimators $\hat{G}^{(k)}$ (defined by (4.1)) obtained from $k$ simulations with the parameters $p \in \mathcal{D}$:

$$E_\omega(p) := \sum_{i \in \mathrm{I}} \omega_i \big\| G_i - \hat{G}_i^{(k)}(p) \big\|_{\ell^2}, \qquad \mathrm{I} = \{sen,\ sen_A,\ sen_B,\ nta\}, \tag{4.2}$$

Similarly, we denote by

$$e_{B\%}(p) := \left| r_{B\%} - \hat{r}_{B\%}^{(k)}(p) \right|, \qquad \hat{r}_{B\%}^{(k)}(p) := \frac{1}{k} \sum_{s=1}^{k} \hat{r}_{B\%}^{s}(p), \tag{4.3}$$

[1]See notations in the first paragraph of Section 4.1. One should not be mistaken: on the "G-graphs" plotted on Fig. 11-*left* the distance is not taken "horizontally" as usual, but "vertically" since the distance is actually between $G^{-1}$ maps.

the error (in absolute value) between experimental and estimated proportions of *type B* lineages among $n_{sen}$ senescent lineages; with $\hat{r}^{(k)}_{B\%}(p)$ the estimator of the proportion defined as the average on $k$ independent simulations with parameters $p$.

The parameters of these error functions are listed in Table 2 and the different parameter domains $\mathcal{D}$ that have been explored are explicitly given in Table 3 of Section 4.4. We denote by $\mathcal{D}_{m,i}$ the $i$th domain of dimension $m$ and by $\mathcal{D}_m$ the union of all the $\mathcal{D}_{m,i}$, $i \geq 1$, introduced.

##### 4.2 Characterization of the estimation method

**CMA-ES estimation strategies and robustness.** Overwhelming empirical evidence agree on CMA-ES convergence for most cost functions, and first theoretical grounds may be found in [6]. To optimize the estimation strategy and reinforce CMA-ES robustness, we followed two classical recommendations:

- Several estimations were run in series: each new simulation kept track of the solutions returned by the previous estimations and started with bigger initial *population sizes* (defined in Table 2). In this case, we denote $N = (N_1, N_2, \ldots, N_l)$ the $l$ successive *population sizes*.

- Every estimation strategy was launched several times in parallel with exact same setting but different *starting point* $p_0$ (again we refer to Table 2) drawn independently in the parameter space. At the end, we checked that most of the solutions obtained were close to each other.

**Sensitivity to the weight $\omega$.** To reinforce the robustness of our results we performed a sensitivity analysis to the weight $\omega$ defining the cost function (4.2), running independent CMA-ES optimizations for various $\omega$. The results have beed reported in [5]. They suggest that the optimization is not very sensitive $\omega$. In the following we chose $\omega := (0.3, 1, 1, 1)$.

##### 4.3 A few comments on the proportion of *type B* cells

Contrarily to the generations of arrest, the proportion of *type B* lineages obtained with our laws ($\hat{r}_{B\%} \approx 74\,\%$) is slightly too high compared with the experimental value ($r_{B\%} \approx 57\,\%$). Although the discrepancy is significant, it must be qualified by the following considerations:

- *The experimental value is slightly biased downwards.* Because of the definition of the classification of lineages, experimental *type A* are necessarily senescent dead lineages ($n_A = n_{sen_A}$), contrarily to *type B* ($n_B = n_{nta} \neq n_{sen_B}$). However not all the $n = 187$ lineages of the dataset have reached senescence: $39 = n - n_{sen}$ have not, because dead accidentally or still alive at the end of the experiment. Among them 31 (=

| | | Parameter |
|---|---|---|
| **CMA-ES** | N | *Population size.* The number of points –in the parameter space $\mathcal{D}$– tested (i.e at which the cost function is evaluated) per iteration, or "generation". At every iteration $i$, the $N$ new points of the $i$th generation are drawn from a normal distribution with parameters $(p_i, \Sigma_i)$ depending on the value of the cost function at the $N$ points of the previous generation. Here $p_i \in \mathcal{D}$ is the approximation of the solution at generation $i$, expected to converge to a minimizer of $E$ while $\Sigma_i$ would get closer to $0_N$. |
| | $p_0$ | *Starting point.* The $N$ points of the generation 0 are drawn from a normal distribution of parameters $(p_0, \sigma_0 D)$, with $D$ is the identity $N$-matrix if $\mathcal{D} = [0,1]^m$, and some rescaled (diagonal) $N$-matrix otherwise. |
| | $\sigma_0$ | *Initial standard deviation.* See above. It should be small if $p_0$ is *a priori* close to the solution. |
| **Error functions** | $\omega$ | *Weights* of the errors from the different graphs, see (4.2) |
| | $\bar{e}_{B\%}$ | *Upper bound for $e_{B\%}$.* For every candidate $p \in \mathcal{D}$, if it is such that the error $e_{B\%}(p)$ (defined by (4.3)) exceeds $\bar{e}_{B\%}$, the value of the cost function at $p$ is set to `NaN` which forces CMA-ES to forget $p$ and draw another point. At the end the optimization is run in the subset of $\mathcal{D}$ where $e_{B\%} \leq \bar{e}_{B\%}$. This is a good way to avoid runtime errors in the simulation of generations arrest of *type A* or B lineages in subsets of $\mathcal{D}$ corresponding to laws of arrest that makes *type A* or B lineages very unlikely (where the proportion of *type B* lineages close to 1 or 0, resp.). |
| | $k$ | *Number of simulations of experimental G-graph* to run and average to approximate the corresponding theoretical graph. |
| | $\|\cdot\|_{\ell^2}$ | *Distance* defining the error between experimental and simulated graphs: $$\|G\|_{\ell^2} := \sqrt{\sum_{l=1}^{n} \left|G(l)\right|^2}, \qquad G \in \mathbb{R}^n$$ NB: we have $n_{nta} = 115$, $n_{sen} = 148$, $n_{sen_A} = 64$ and $n_{sen_B} = 84$. |

Table 2: Table of the parameters of the optimization method, distinguishing between parameters of the optimizer (CMA-ES) and of the error functions ($E$ and $e_{B\%}$).

$n_B - n_{sen_B}$) identify as *type B*, while only $n_X = 8$ are unclassified. As a result, there are proportionally more *type B* lineages than *type A* in the whole dataset than in the subdataset of senescent lineages:

$$\bar{r}_{B\%} \approx 100 \times \frac{n_B}{n_A + n_B} \approx 64\,\%, \qquad \bar{r}_{B\%} \in \left[\bar{r}_{low}, \bar{r}_{up}\right],$$
$$\bar{r}_{low} := 100 \times \frac{n_B}{n_B + n_A + n_X} \approx 61\,\%, \qquad \bar{r}_{up} := 100 \times \frac{n_B + n_X}{n_B + n_A + n_X} \approx 67\,\%,$$

with $\bar{r}_{B\%}$ be the proportion of type B lineages in the whole dataset.

- *The confidence interval is sufficiently large.* The 5$^{\text{th}}$ and 95$^{\text{th}}$ percentiles of a $k$-sample, for $k = 1000$ simulations, of the proportion $\hat{r}_{B\%}$ given by our laws are around 67 % and 80 %, and the extrema around 61 % and 89 %.

These two points reconcile the discrepancy between the experimental and simulated proportions of *type B* lineages, since $\bar{r}_{B\%}$ belong to the (empirical) 90 % confidence interval given by our laws.

##### 4.4 More details on the estimations carried out

The parameter estimation constitutes one of our main result: when senescence onset is assumed to be stochastically driven by the shortest telomere, experimental data are best fitted by senescence laws that are qualitatively very distinct for *type A* and *type B* cells.

The strength of such result relies on the estimation procedure, that we expose here. In particular, we first assumed that $p_{sen_A}$ and $p_{sen_B}$ were identical (equal to $p_{sen}$), i.e. that senescence is triggered similarly for each type. This assumption however is not sufficient to fit the data (Fig. 13 and Fig. 14) and we authorized $p_{sen_A}$ and $p_{sen_B}$ to be different. The obtained result is striking since experimental data are then very well fitted and the estimated laws $p_{sen_A}$ and $p_{sen_B}$ are not close at all (Fig. 19).

Let us recall the probability laws whose parameters were estimated.

- The probability for the shortest telomere of length $\ell$ to trigger a first sequence of non-terminal arrest(s):

$$p_{nta}(\ell) = \min\big(1, b_{nta}\,\mathrm{e}^{-a_{nta}\ell}\big), \quad (a_{nta}, b_{nta}) \in (0,1]^2. \tag{$\mathcal{P}_{nta}$}$$

- The probability for a cell with shortest telomere of length $\ell$ to enter senescence when it is assumed independent of cell type: for $(a_{sen}, b_{sen}, \ell_{min}) \in [0,1] \times (0,1] \times \mathbb{Z}$,

$$p_{sen}(\ell) = \begin{cases} \min\big(1, b_{sen}\,\mathrm{e}^{-a_{sen}\ell}\big), & \text{if } \ell > \ell_{min}, \\ 1, & \text{if } \ell \leq \ell_{min}. \end{cases} \tag{$\mathcal{P}_{sen}$}$$

- The previous law when we allow it to be different for *type A* and *type B* cells:

$$p_{sen_A}(\ell) = \begin{cases} \min\big(1, b_{sen_A}\,\mathrm{e}^{-a_{sen_A}\ell}\big), & \text{if } \ell > \ell_{min_A}, \\ 1, & \text{if } \ell \leq \ell_{min_A}, \end{cases} \tag{$\mathcal{P}_{sen_A}$}$$

$$p_{sen_B}(\ell) = \begin{cases} \min\big(1, b_{sen_B}\,\mathrm{e}^{-a_{sen_B}\ell}\big), & \text{if } \ell > \ell_{min_B}, \\ 1, & \text{if } \ell \leq \ell_{min_B}. \end{cases} \tag{$\mathcal{P}_{sen_B}$}$$

In addition, we fitted the parameters $\ell_{trans}$, $\ell_0$ and $\ell_1$ of the initial distribution of telomere lengths $f_{init}$. Their corresponds to specific transformations (see equation (1.1)) of the initial distribution $f_0$ of Bourgeron et al. [3].

The different domains on which estimations have been run are described on Table 3. We denote by $\mathcal{D}_{m,i}$ the $i$th domain of dimension $m$ and by $\mathcal{D}_m$ the union of all the $\mathcal{D}_{m,i}$, $i \geq 1$, introduced.

As explained in Section 4.2, we ran, for every parameter domain, several independant CMA-ES estimations so that to improve the robustness of the results. Each estimation gave us a point in the parameter domain, i.e. a set of parameters to which correspond spefic laws estimated to best minimize the cost function (4.2). For each parameter domain, the set of these estimated best-fit laws are plotted on Fig. 12, 15 and 18. We see that they are qualitatively close which reinforce the idea that the minimum approached through our estimation procedure is global. In other words, it is unlikely that there exist qualitatively different laws allowing to fit the experimental data as well.

| Domain | Parameter range | | | | | |
|---|---|---|---|---|---|---|
| | $(\mathcal{P}_{nta})$ | | $(\mathcal{P}_{sen})$ | | | $f_{init}$ |
| | $a_{nta}$ | $b_{nta}$ | $a_{sen}$ | $b_{sen}$ | $\ell_{min}$ | $\ell_0$ |
| $\mathcal{D}_{5,1}$ | $[0,1]$ | $[0,1]$ | $[0,1]$ | $[0,2]$ | $[\![0,80]\!]$ | $\{0\}$ |
| $\mathcal{D}_{5,2}$ | $[0,1]$ | $[0,1]$ | $[0,1]$ | $[0,1]$ | $[\![20,80]\!]$ | $\{0\}$ |
| $\mathcal{D}_{6}$ | $[0,1]$ | $[0,1]$ | $[0,1]$ | $[0,1]$ | $[\![27,40]\!]$ | $[\![0,30]\!]$ |

(a) Senescence entry assumed identical for *type A* and *type B* cells

| Domain | Parameter range | | | | | | | | | | |
|---|---|---|---|---|---|---|---|---|---|---|---|
| | $(\mathcal{P}_{nta})$ | | $(\mathcal{P}_{sen_A})$ | | | $(\mathcal{P}_{sen_B})$ | | | $f_{init}$ | | |
| | $a_{nta}$ | $b_{nta}$ | $a_{sen_A}$ | $b_{sen_A}$ | $\ell_{min_A}$ | $a_{sen_B}$ | $b_{sen_B}$ | $\ell_{min_B}$ | $\ell_{trans}$ | $\ell_0$ | $\ell_1$ |
| $\mathcal{D}_{7}$ | $[0,1]$ | $[0,1]$ | $[0,1]$ | $[0,1]$ | $\{30\}$ | $[0,1]$ | | $[\![0,40]\!]$ | | | |
| $\mathcal{D}_{8,1}$ | $[0,1]$ | $[0,1]$ | $[0,1]$ | $[0,2]$ | $[\![0,80]\!]$ | $[0,1]$ | $[0,2]$ | $[\![0,80]\!]$ | $\{0\}$ | | |
| $\mathcal{D}_{8,2}$ | $[0,1]$ | $[0,1]$ | $[0,1]$ | $[0,1]$ | $[\![20,80]\!]$ | $[0,1]$ | $[0,1]$ | $[\![0,60]\!]$ | | $\{0\}$ | $\{0\}$ |
| $\mathcal{D}_{8,3}$ | $[0,1]$ | $[0,1]$ | $[0,1]$ | $[0,1]$ | $[\![27,40]\!]$ | $[0,1]$ | $[0,1]$ | $[\![0,40]\!]$ | | | |
| $\mathcal{D}_{9,1}$ | $[0,0.1]$ | $[0.1,0.8]$ | $[0,1]$ | $[0,1]$ | $[\![27,40]\!]$ | $[0,1]$ | $[0,1]$ | $[\![0,40]\!]$ | $[\![0,30]\!]$ | | |
| $\mathcal{D}_{9,2}$ | $[0,1]$ | $[0,1]$ | $[0,1]$ | $[0,1]$ | $[\![27,40]\!]$ | $[0,1]$ | $[0,1]$ | $[\![0,40]\!]$ | $[\![0,30]\!]$ | | |
| $\mathcal{D}_{10,1}$ | $[0,0.1]$ | $[0.1,0.8]$ | $[0,1]$ | $[0,1]$ | $[\![27,40]\!]$ | $[0,1]$ | $[0,1]$ | $[\![0,40]\!]$ | 0 | $[\![20,30]\!]$ | $[\![0,60]\!]$ |
| $\mathcal{D}_{10,2}$ | $[0,1]$ | $[0,1]$ | $[0,1]$ | $[0,1]$ | $[\![0,40]\!]$ | $[0,1]$ | $[0,1]$ | $[\![0,30]\!]$ | 0 | $[\![0,30]\!]$ | $[\![0,60]\!]$ |

(b) Senescence entry assumed different for *type A* and *type B* cells

Table 3: Tables of the domains in which parameter estimations were run.

##### Same senescence law for *type A* and *type B* cells

We first estimated the best parameters of $(\mathcal{P}_{nta})$ and $(\mathcal{P}_{sen})$ when senescence entry is assumed statistically identical for *type A* and *type B* cells. Different domains were successively tested:

1. $\mathcal{D}_{5,1}$: the distribution $f_{init}$ of initial telomere lengths is fixed (equal to $f_0$, i.e. $(\ell_{trans}, \ell_0, \ell_1) = (0, 0, 0)$) and $\ell_{min}$ can be 0 bp,

2. $\mathcal{D}_{5,2}$: same as $\mathcal{D}_{5,1}$ except than $\ell_{min}$ is forced to be more than 20 bp,

3. $\mathcal{D}_6$: $\ell_{min}$ is more than 27 bp but $f_{init} = f_{init}(\cdot; \ell_{trans}, 0, 0)$ is a translation of $f_0$ by a value $\ell_T$ that is fitted.

On each domain, the results of the independent parameter estimations that were run are gathered in Fig. 12. They are qualitatively close to each other which is a good indicator that the estimations have converged close to a global minimum. The best result among all these independent estimates (Fig. 13) was then used for simulation. The resulting generation graphs are compared to experimental ones on Fig. 14.

Under the assumption that the law of senescence onset is identical for *type A* and *type B*, and providing that we either let $\ell_{min}$ be close to 0 ($\mathcal{D}_{5,1}$) or start from a translated distribution of telomere lengths ($\mathcal{D}_6$), we see that our model approaches well all experimental graphs except $G_{sen_B}$ which lacks variability. Such a discrepancy reinforce the idea that the law of senescence entry could be different depending on cells type.

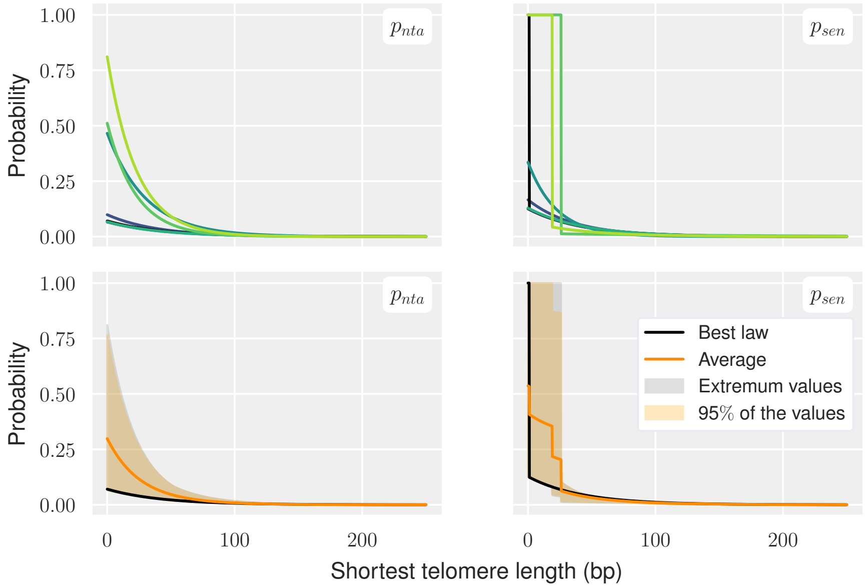


(a) $\mathcal{D} = \mathcal{D}_{5,1}$

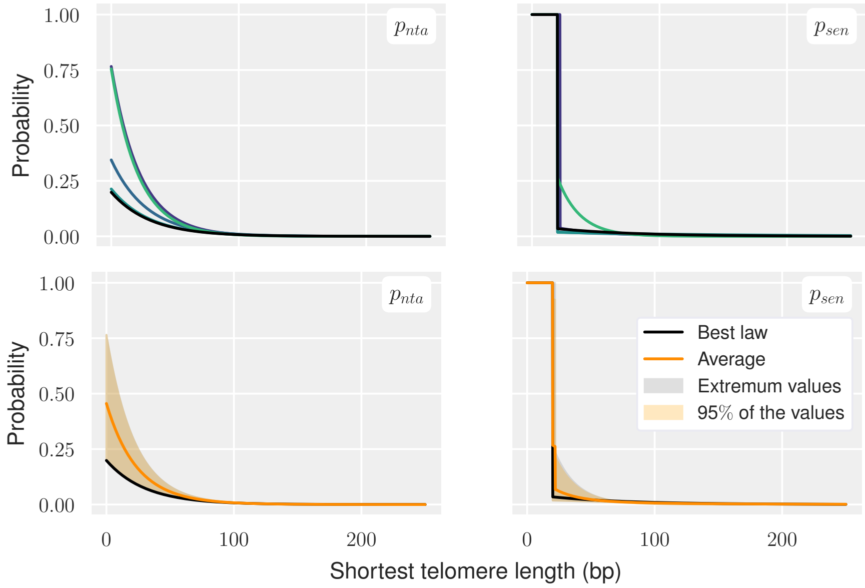


(b) $\mathcal{D} = \mathcal{D}_{5,2}$

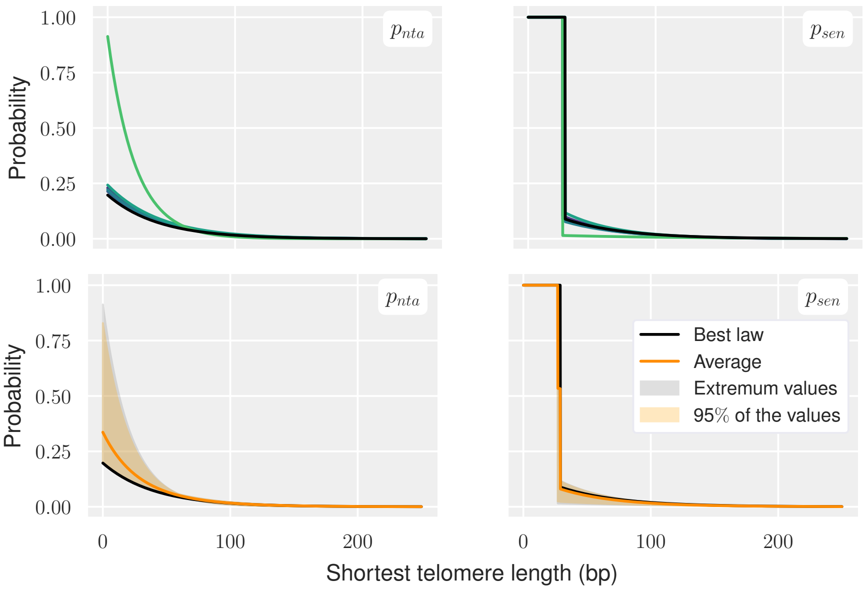


(c) $\mathcal{D} = \mathcal{D}_6$

Figure 12: Estimates of the laws $(\mathcal{P}_{nta})$, $(\mathcal{P}_{sen})$ retrieved when running several independent CMA-ES estimations on different domains $\mathcal{D}$. Two different representations: *(top)* either plotting each fit individually, or the average fit with $5th$ and $95th$ percentile *(bottom)*. The black line *(bottom)* corresponds to the fit that best minimizes $E_{\omega^2}$, plotted on Fig. 13.

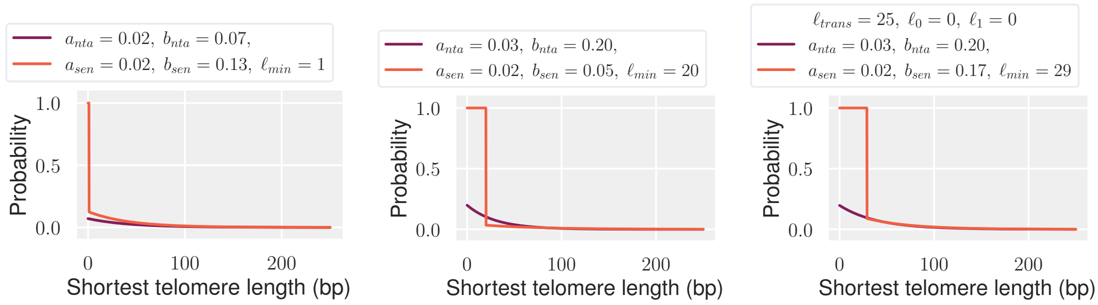


(a) $\mathcal{D}_{5,1}$: $f_{init} = f_0$ fixed, $\ell_{min} \geq 0$

(b) $\mathcal{D}_{5,2}$: $f_{init} = f_0$ fixed, $\ell_{min} \geq 20$

(c) $\mathcal{D}_6$: $f_{init} = f_0(\cdot - \ell_T)$ fitted through $\ell_T$, $\ell_{min} \geq 27$

Figure 13: Best fit of the laws $(\mathcal{P}_{nta})$ and $(\mathcal{P}_{sen})$ retrieved by CMA-ES estimations run on the different domains $\mathcal{D}_{5,1}$, $\mathcal{D}_{5,2}$ and $\mathcal{D}_6$.

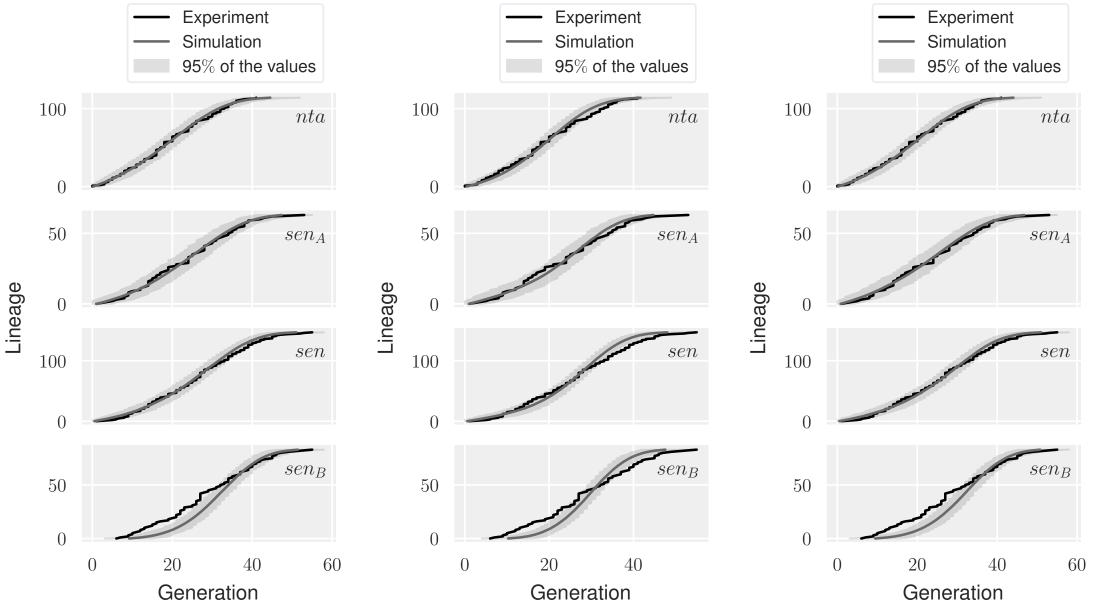


(a) $\mathcal{D}_{5,1}$: $f_{init} = f_0$ fixed, $\ell_{min} \geq 0$

(b) $\mathcal{D}_{5,2}$: $f_{init} = f_0$ fixed, $\ell_{min} \geq 20$

(c) $\mathcal{D}_6$: $f_{init} = f_0(\cdot - \ell_T)$ fitted through $\ell_T$, $\ell_{min} \geq 27$

Figure 14: Comparison to experimental data of the generation curves simulated with the best fits of the laws $(\mathcal{P}_{nta})$ and $(\mathcal{P}_{sen})$ plotted on Fig. 13. Average on $k = 1000$ simulations, $5^{\text{th}}$ and $95^{\text{th}}$ percentiles and extremum values.

##### Different senescence law for *type A* and *type B* cells

*Allowing translations of $f_0$.* Following the same method, we estimated the 8 parameters of the laws $(\mathcal{P}_{nta})$, $(\mathcal{P}_{sen_A})$ and $(\mathcal{P}_{sen_B})$, on different domains, possibly allowing translations of $f_0$ as well:

1. $\mathcal{D}_{8,1}$: $f_{init} = f_0$ is not fitted and $\ell_{min_A}$ can be 0 bp,
2. $\mathcal{D}_{8,2}$: same as $\mathcal{D}_{8,1}$ except than $\ell_{min_A}$ is forced to be more than 20 bp,
3. $\mathcal{D}_9$: $\ell_{min_A}$ is more than 27 bp but $f_{init} = f_{init}(\cdot; \ell_{trans}, 0, 0)$ is fitted through $\ell_{trans}$.

The results, plotted on Fig. 15 to 17 are very similar to those derived previously (see Fig. 12 to 14 respectively) in the sense that to have, at least for *type A* cells, a threshold $\ell_{min_A} \geq 20$ bp requires to translate the initial distribution of telomere lengths of about $\ell_{min_A}$. The novelty is that all the graphs are now perfectly fitted (Fig. 17).

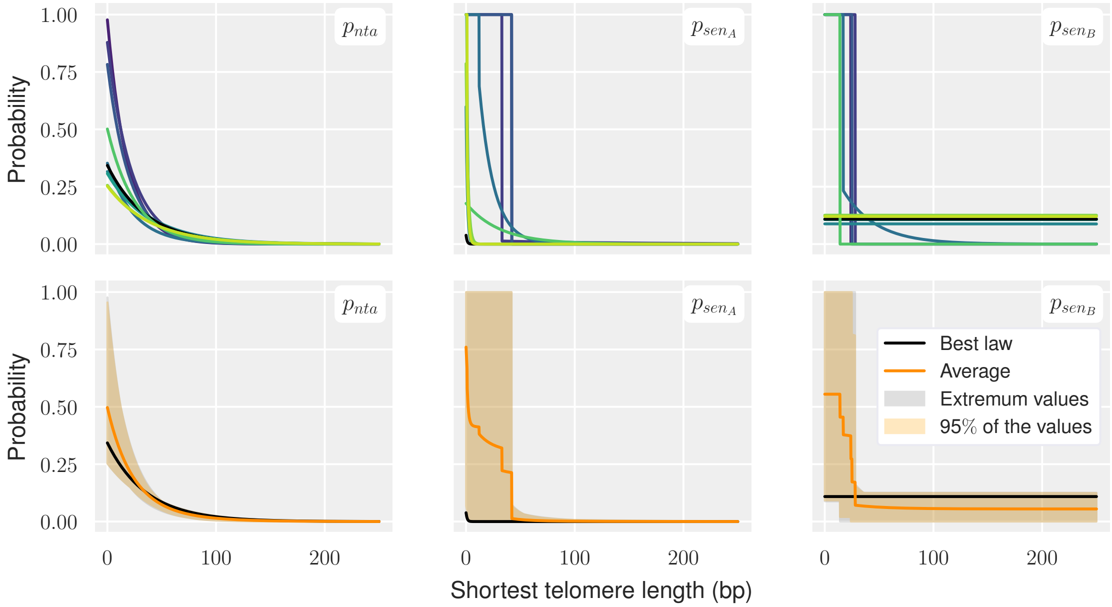


(a) $\mathcal{D} = \mathcal{D}_{8,1}$

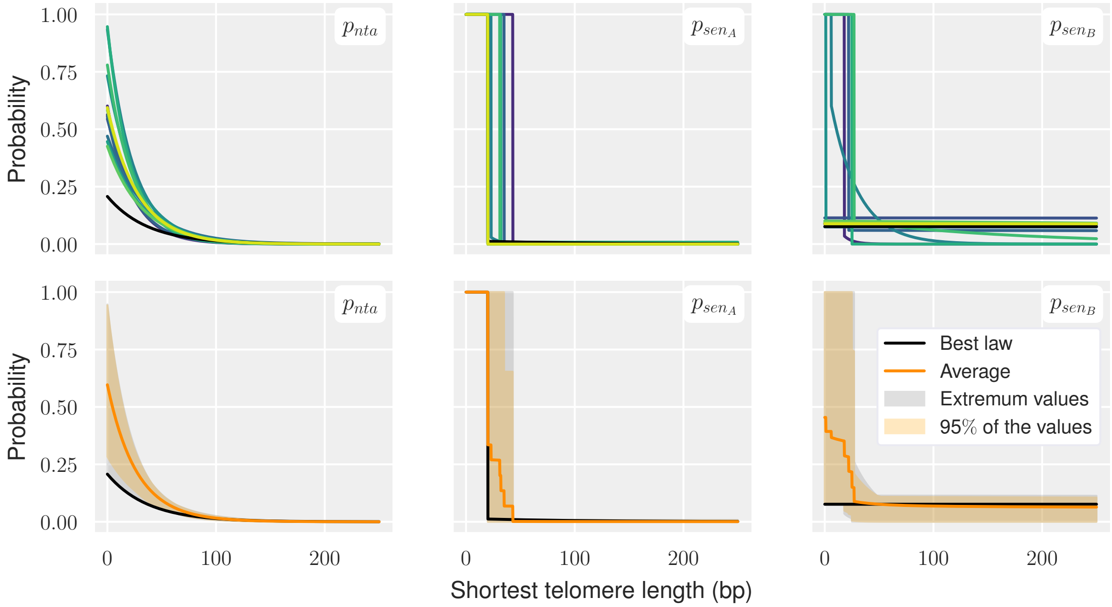


(b) $\mathcal{D} = \mathcal{D}_{8,2}$

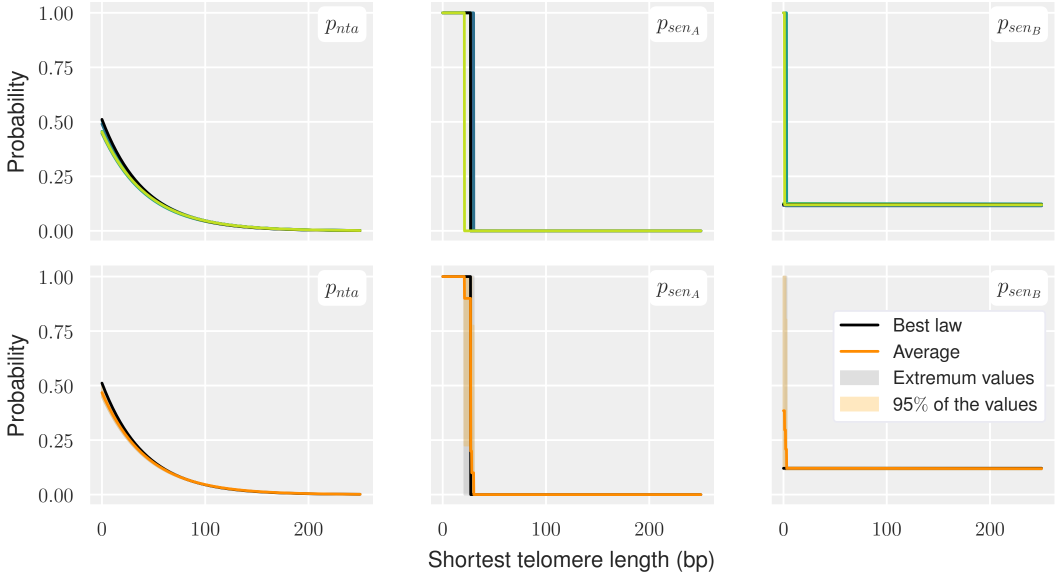


(c) $\mathcal{D} = \mathcal{D}_9$

Figure 15: Estimates of laws $(\mathcal{P}_{nta})$, $(\mathcal{P}_{sen_A})$ and $(\mathcal{P}_{sen_B})$ retrieved when running several independent CMA-ES estimations on different domains $\mathcal{D}$. Two different representations: *(top)* either plotting each fit individually, or the average fit with $5th$ and $95th$ percentile *(bottom)*. The black line *(bottom)* corresponds to the fit that best minimizes $E_{\omega^3}$, plotted on Fig. 16.

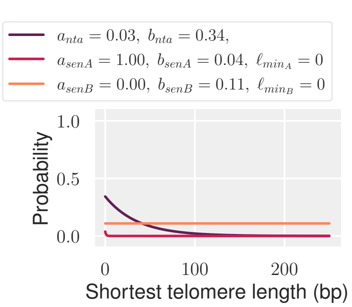


(a) $\mathcal{D}_{8,1}$: $f_{init} = f_0$ fixed, $\ell_{min_A} \geq 0$

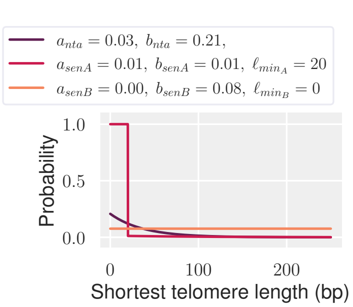


(b) $\mathcal{D}_{8,2}$: $f_{init} = f_0$ fixed, $\ell_{min_A} \geq 20$

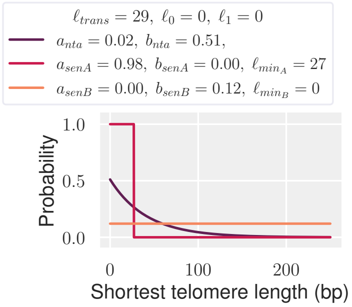


(c) $\mathcal{D}_9$: $f_{init} = f_0(\cdot - \ell_T)$ fitted through $\ell_T$, $\ell_{min_A} \geq 27$

Figure 16: Best fit of the laws $(\mathcal{P}_{nta})$, $(\mathcal{P}_{sen_A})$ and $(\mathcal{P}_{sen_B})$ retrieved by CMA-ES estimations run on the different domains $\mathcal{D}_{8,1}$, $\mathcal{D}_{8,2}$ and $\mathcal{D}_9$.

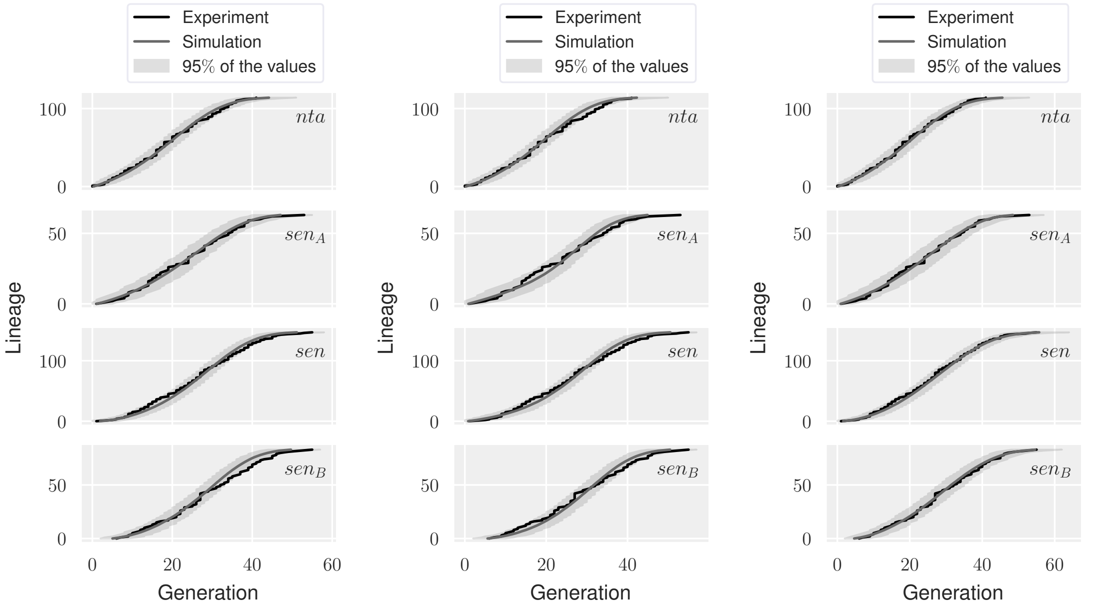


(a) $\mathcal{D}_{8,1}$: $f_{init} = f_0$ fixed, $\ell_{min_A} \geq 0$

(b) $\mathcal{D}_{8,2}$: $f_{init} = f_0$ fixed, $\ell_{min_A} \geq 20$

(c) $\mathcal{D}_9$: $f_{init} = f_0(\cdot - \ell_T)$ fitted through $\ell_T$, $\ell_{min_A} \geq 27$

Figure 17: Comparison to experimental data of the generation curves simulated with the best fits of the laws $(\mathcal{P}_{nta})$, $(\mathcal{P}_{sen_A})$ and $(\mathcal{P}_{sen_B})$ plotted on Fig. 16. Average, 5$^{\text{th}}$ and 95$^{\text{th}}$ percentiles and extremum values on $k = 1000$ simulations.

*Allowing dilatation of* $f_0$. What has probably proved useful in the previous estimations to account for non-zero $\ell_{min}$ or $\ell_{min_A}$ –that we think more biologically relevant than close-to-zero thresholds– is likely the translation of the infimum $\ell_{inf}$ of the support of $f_0$ specifically, rather than the translation of the whole distribution $f_0$.

To achieve the same goal while preserving the mode of $f_0$ (not preserved by translations) we eventually fitted $\ell_0$ and $\ell_1$ together with the parameters of the laws $(\mathcal{P}_{nta})$, $(\mathcal{P}_{sen_A})$ and $(\mathcal{P}_{sen_B})$.

All estimates were very close (Fig. 18). The best fit retrieved (Fig. 19b), is the one we decided to choose for its excellent accordance with experimental data (Fig. 19a).

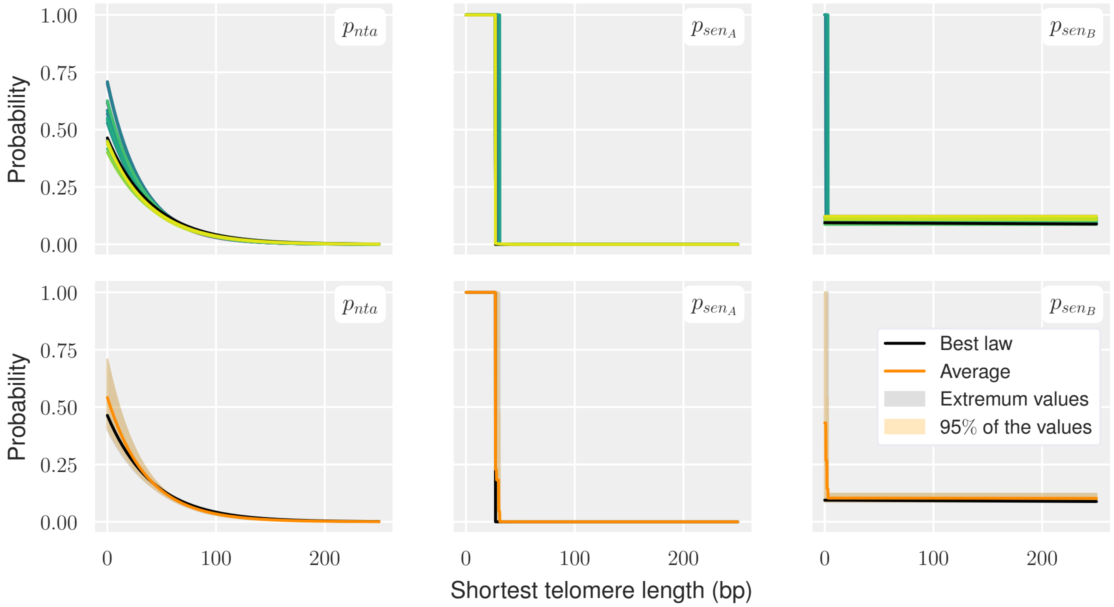


Figure 18: Estimates of the laws $(\mathcal{P}_{nta})$, $(\mathcal{P}_{sen_A})$ and $(\mathcal{P}_{sen_B})$ retrieved when running several independent CMA-ES estimations on $\mathcal{D}_{10}$. Two different representations: *(top)* either plotting each fit individually, or the average fit with $5th$ and $95th$ percentile *(bottom)*. The black line *(bottom)* corresponds to the fit that best minimizes $E_{\omega^3}$.

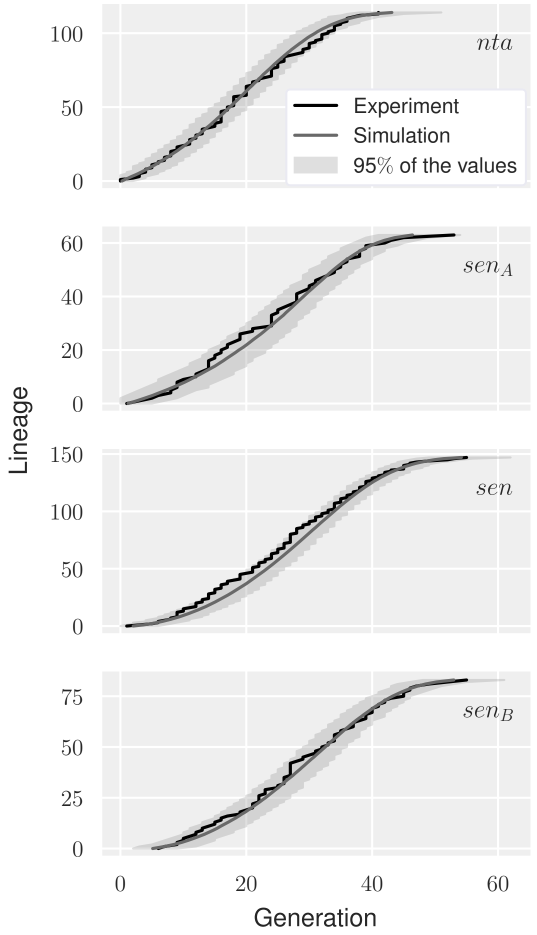


(a) Comparison of the simulated generation curves to experimental data

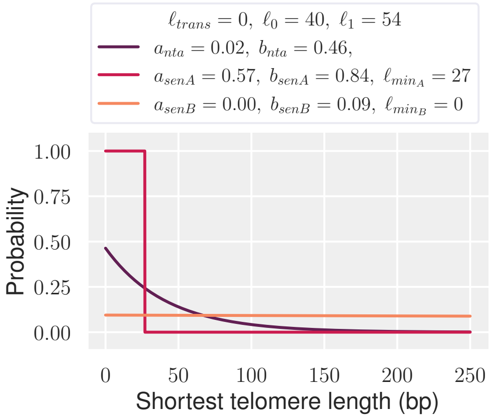


(b) Best fits of the laws $(\mathcal{P}_{nta})$, $(\mathcal{P}_{sen_A})$ and $(\mathcal{P}_{sen_B})$ and the dilatation parameters $\ell_0$ and $\ell_1$

Figure 19: Results of the estimation of the laws $(\mathcal{P}_{nta})$, $(\mathcal{P}_{sen_A})$ and $(\mathcal{P}_{sen_B})$ run in $\mathcal{D}_{10}$ defined on Table 3b.

[5] Anais Rat. *Structured population dynamics: theory, asymtotic and numerical analysis. Application to populations heterogeneous in growth rate and to the replicative senescence.* PhD thesis, Ecole Centrale Marseille, 2023.

[6] Cheikh Touré, Anne Auger, and Nikolaus Hansen. Global linear convergence of evolution strategies with recombination on scaling-invariant functions. *Journal of Global Optimization*, 86(1):163–203, 2023.

[7] Zhou Xu, Khanh Dao Duc, David Holcman, and Maria Teresa Teixeira. The length of the shortest telomere as the major determinant of the onset of replicative senescence. *Genetics*, 194(4):847–857, 2013.

[8] Zhou Xu, Emilie Fallet, Camille Paoletti, Steffen Fehrmann, Gilles Charvin, and Maria Teresa Teixeira. Two routes to senescence revealed by real-time analysis of telomerase-negative single lineages. *Nature Communications*, 6(1), 2015.
